# Biomineralization and biomechanical trade-offs under heterogeneous environments in the eastern oyster *Crassostrea virginica*

**DOI:** 10.1101/2023.11.28.569049

**Authors:** Luca Telesca, Braddock K. Linsley, Lukasz Witek, Bärbel Hönisch

## Abstract

Accurate biological models are critical to reliably predict vulnerability of marine organisms and ecosystems to rapid environmental changes. Current predictions on the biological impacts of climate change and human-caused disturbances primarily stem from controlled experiments but lack assessments of the mechanisms underlying biotic variations in natural systems. Such information is key to translating experimental models to natural populations, especially for habitat-forming, climate sensitive species with key ecological roles. This study aimed to characterize and quantify spatial patterns of shell biomineralization and biomechanical properties in a key reef-building oyster, *Crassostrea virginica*, collected from restored reefs along natural estuarine gradients in the Hudson River Estuary (NY, U.S.). We characterized patterns of oyster shell production (i.e., shape and thickness), structure (i.e., abundance of foliated and chalky calcite), mineralogy (i.e., crystal size and density), composition (i.e., organic matrix and Mg/Ca ratios), and mechanical performance (i.e., elastic modulus and hardness) at the macro and micro scale. Our results demonstrate a strong protective capacity of *C. virginica* for compensatory adjustments in shell biomineralization and biomechanics to maintain shell production and protective functions as a response to biotic and abiotic stressors. We reveal salinity as a key predictor of oyster shell structure, mechanical integrity, and resistance to dissolution, and describe the functional role of chalky calcite in shaping shell mechanical performance. Compensatory adjustments along salinity gradients indicate that oysters produce shells with *i*) high mechanical resistance but increased vulnerability to dissolution under marine conditions, and *ii*) lower structural integrity but higher protection from dissolution under brackish conditions. Our work illustrates that biomineralization and biomechanical adjustments may act as compensatory mechanisms in eastern oysters to maintain overall performance under heterogeneous estuarine environments, and could represent a cornerstone for calcifying organisms to acclimate and maintain their ecological functions in a rapidly changing climate.

## INTRODUCTION

Although the ocean moderates anthropogenic climate change, this has significant impacts on its fundamental physics and chemistry (Gattuso *et al*., 2015; IPCC, 2022), with dramatic effects on marine organisms and ecosystem functions (Nagelkerken & Connell, 2015; Ellis, Urbina & Wilson, 2017). While current predictions on the biological impacts of climate and human-caused disturbances mostly stem from controlled experiments (Wernberg, Smale & Thomsen, 2012; Urban *et al*., 2016), they generally lack assessments of the mechanisms shaping biotic variations in natural environments (Capotondi *et al*., 2019). Yet, this information is critical to translate experimental models to natural populations (Connell *et al*., 2017; Vargas *et al*., 2017; Telesca *et al*., 2019), especially for habitat-forming species with high climate sensitivity, such as calcifying organisms.

Marine calcifiers, which produce calcium carbonate (CaCO_3_) shells and skeletons, are often key to maintaining healthy, functioning marine and estuarine ecosystems (Gutierrez *et al*., 2003), but are predicted to face significant challenges under rapidly changing climates (Figuerola *et al*., 2021). In recent years, however, studies have shown some marine calcifiers to exhibit compensatory mechanisms that can counteract or even reverse the impacts of environmental change on overall species performance (Connell *et al*., 2017; Leung, Russell & Connell, 2017; Cross, Harper & Peck, 2019; Telesca *et al*., 2021). These mechanisms act through a range of morphological and physiological responses of calcification, including adjustments of skeletal growth (Leung, Russell & Connell, 2017), deposition (Cross, Harper & Peck, 2019; Telesca *et al*., 2021), mineralogy (Telesca *et al*., 2019), mechanical properties (Meng, Fitzer *et al*., 2018) and organic content (Lagos *et al*., 2021).

This body of research has primarily addressed the influence of ocean warming and acidification, but studies have rarely considered the effect of other environmental changes, such as salinity gradients (Durack, Wijffels & Matear, 2012). Salinity is a leading physical driver of biological adaption (R. C. Newell, 1976) and ecosystem functioning (Smyth & Elliott, 2016), as well as species interactions (Kimbro *et al*., 2017) and bio-calcification (Thomsen, *et al*., 2015). Indeed, salinity positively correlates with increasing predators’ abundance (e.g., crabs, star fish, drilling gastropods, and fish) and increasing predation pressure on marine calcifiers (Kimbro *et al*., 2017; Pusack *et al*., 2019; Dickey *et al*., 2021). Moreover, the availability of the chemical building blocks of calcification (i.e., bicarbonate [HCO_3_ ^−^] and calcium [Ca^2+^] ions) decreases with seawater salinity, influencing the stability of CaCO_3_ structures through *i*) a decreased CaCO_3_ saturation state (Ω_CaCO3_) (Thomsen *et al*., 2018), *ii*) unfavorable ratios of calcification substrates to the inhibitor [H^+^] (Sanders *et al*., 2021), and *iii*) higher calcification costs (Sanders *et al*., 2018). Saturation state of CaCO_3_ polymorphs (predominantly calcite or aragonite) controls calcification kinetics (Waldbusser *et al*., 2015) and net deposition or dissolution of biogenic CaCO_3_ (Ries *et al*., 2016). In the coastal region and estuaries, salinity is highly variable and strongly influenced by precipitation and terrestrial run-off (Röthig *et al*., 2023). This variability has been further exacerbated by the stronger anthropogenic impacts on biogeochemical cycles in nearshore environments compared to the open ocean (Durack, Wijffels & Matear, 2012; Levin *et al*., 2015). Aside from the effects on ocean physical processes, the impacts of variable salinity on marine calcifiers’ structural integrity and ecological functions are largely unknown (Telesca *et al*., 2019).

The eastern oyster, *Crassostrea virginica* (Gmelin, 1791), is a reef-forming bivalve with a significant ecological, cultural and economic value in most estuaries along the North American eastern margin (MacKenzie, 1996; Byers *et al*., 2006; FAO, 2018). Eastern oyster reefs enhance estuary biodiversity and offer valuable ecosystem services, including estuarine integrity and resilience (Byers *et al*., 2006; Hastings *et al*., 2007), fish nurseries (Grabowski & Peterson, 2007), biofiltration (R. I. E. Newell, 2004), and coastal protection (Grabowski & Peterson, 2007; Rodriguez *et al*., 2014). However, habitat degradation, overfishing, and environmental change have decimated *C. virginica* reefs’ dramatically during the last century (Beck *et al*., 2011; Zu Ermgassen *et al*., 2012), making this bivalve a focus of intensive restoration efforts (Beck et al., 2011).

Oyster shell integrity provides vital structural support along with protection against predators and corrosive water conditions, making shell traits subject to strong selective pressure (John D. Taylor & Layman, 1972; Lombardi *et al*., 2013). Eastern oyster shells are composed of calcitic crystals, with a minor fraction of organic matrix (up to 6%), organized in three main microstructures: *i*) a thin prismatic layer, *ii*) compact foliated layers, and *iii*) porous chalky deposits (John David Taylor, Kennedy & Hall, 1973). These microstructures are characterized by differences in mechanical performance, solubility, and energetic costs (A. R. Palmer, 1992; Harper, 2000; S. W. Lee, Kim & Choi, 2008; S.-W. Lee *et al*., 2011), the combination of which determines shells’ overall mechanical and chemical protection (Meng, Fitzer *et al*., 2018; Chadwick *et al*., 2019). Experimental studies have shown unequivocal impacts of controlled physical, chemical and biological changes on oyster shell integrity (Waldbusser *et al*., 2011; Meng, Guo *et al*., 2018; Schwaner *et al*., 2023). But despite projected alteration of estuarine salinity (Röthig *et al*., 2023) and, therefore, predation pressure (Kimbro *et al*., 2017), water carbonate chemistry and calcification costs (Sanders *et al*., 2018; Thomsen *et al*., 2018), the influence of salinity gradients on oyster shell structure and properties is poorly understood (Dickinson *et al*., 2012).

In this study, we characterize and quantify shell biomineralization and biomechanical responses in wild *Crassostrea virginica* specimens collected from restored reefs in the Hudson River Estuary (NY, U.S.) along natural estuarine gradients. Given the capacity for phenotypic plasticity in shell morphology of oysters in response to altered environments and predation pressure (Lord & Whitlatch, 2012; Scherer & Smee, 2017; Meng, Fitzer *et al*., 2018), we test the hypothesis that *C. virginica* can exhibit compensatory responses to maintain shell integrity and functions in response to environmental stress by measuring macroscale and microscale shell structure, composition, and mechanical performance. If *C. virginica* has the capacity for biomineralization and biomechanical adjustments as compensatory responses to changing growth conditions, it might be a cornerstone to understanding acclimation mechanisms in highly dynamic estuarine environments to maintain overall performance and ecological functions, along with reefs’ ecosystem services, under rapidly changing climates.

## MATERIAL AND METHODS

### Oyster collection

Between November 2020 and January 2021, we collected *Crassostrea virginica* individuals from four restored reefs across the Hudson River Estuary (HRE) (Fig. 1). A total of 75 adult oysters of various sizes (shell height 60-180 mm) were collected from four subtidal reefs (depth of ∼0.5 m) as part of the Billion Oyster Project (BOP) seasonal monitoring program (Fig. 1; for details, see Supplementary Material Table S1). For each specimen, we removed the soft tissue, measured the shell weight, and acquired digital photographs to measure shell area (as a proxy for size).

**Figure 1.**
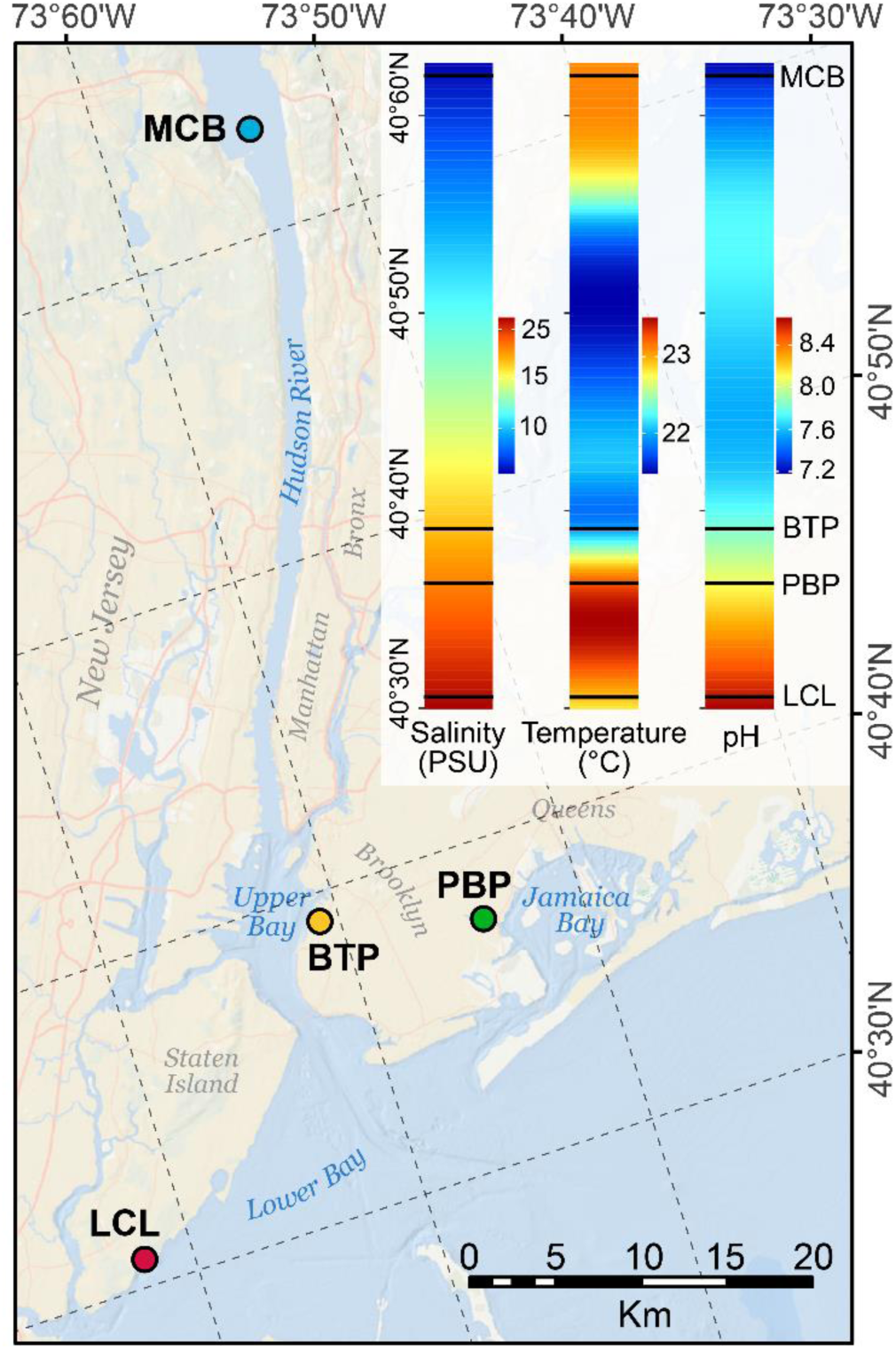
Map of the HRE system in the NY state showing locations of the sampled restored reefs at (*i*) the Governor Mario M. Cuomo Bridge (MCB, *n* = 18) in the Lower Hudson River, (*ii*) the Bush Terminal Piers Park (BTP, *n* = 18) in the Upper Bay, (*iii*) the Paerdegat Basin Park (PBP, *n* = 20) in the Jamaica Bay, and (*iv*) the Lemon Creek Lagoon (LCL, *n* = 19) in the Lower Bay. Conceptual heat maps illustrate the covariation of multiple water parameters across the region and study locations (horizontal black lines). Heat maps were interpolated using average values from June to October 2017-2020 monitoring data at the study sites and HRECOS data. Map created with ArcMap (ArcGIS v10.5 by Esri), background image courtesy of Esri Ocean Basemap (Sources: Esri, Garmin, GEBCO, NOAA NGDC, and other contributors).

These locations were selected because: (*i*) they are representative of the marked North-South environmental gradients in the HRE, especially salinity (7-28 PSU), (*ii*) long-term water-quality monitoring has been implemented by the BOP since 2016 (Baumann, Burmester & Castro, 2021) and the Hudson River Environmental Conditions Observing System (HRECOS; www.hrecos.org) since 2010, (*iii*) the selected reefs were installed at the same time (June-August 2018; Supplementary Material Table S1) using similar restoration substrates (i.e., metal gabions), and (*iv*) reefs were characterized by recruitment (new spat) since first install (Baumann, Burmester & Castro, 2021; Lodge *et al*., 2021).

### Environmental datasets

To assess environmental effects on oyster biomineralization, we gathered available datasets of water temperature, salinity, pH, turbidity, dissolved oxygen (DO) and chlorophyll-*a* (Chl-*a*) concentration, which serves as a proxy for food supply (C. R. Newell *et al*., 2021), for each location. We selected these parameters based on their known influences on oyster growth (C. R. Newell *et al*., 2021) and their sensitivity to climate change, as projected by the IPCC 2022 (IPCC, 2022) report. Water quality data, including continuous and point measurements, at the LCL, BTP, and PBP locations were collected between June and October (the oyster growth season in HRE) from 2017 to 2020 and made available by the BOP (Burmester & McCann, 2018; Baumann, Burmester & Castro, 2021) (details in Supplementary Document S1). For location MCB, continuous water quality data were obtained from the HRECOS station on the Piermont Pier (NY), which is adjacent to the MCB site (details in Supplementary Document S1).

### Macroscale and Microscale analyses

We measured macroscale and microscale properties of shells from each of the four study locations. Macroscale traits, including shell outline shape, thickness, structure, flexural properties (elastic modulus; increasing modulus = brittle shell), density, and bulk organics content, were measured on all specimens (*n* = 75). A subset of five specimens for each location (*n* = 20) was used to quantify microscale shell traits, including organic matrix content, crystal traits, Mg/Ca ratios, calcitic crystals’ elastic modulus and hardness.

### Elliptic Fourier analysis of shell outlines

To quantify shell shapes, we employed a geometric morphometrics approach, which was based on an Elliptic Fourier Analysis (EFA) of outlines (Kuhl & Giardina, 1982). Outlines of lateral and ventral shell views were digitized, processed, and analysed with an EFA as described by Telesca *et al*. (2018) (see Supplementary Methods S1 and Supplementary Material Fig. S1-2) using the Momocs package (Bonhomme *et al*., 2014) in the software R (v4.2.2) (R Core Team, 2022). For each shell outline, four coefficients per calculated harmonic (28 descriptors) were extracted and used as variables quantifying geometric information. A principal component analysis (PCA) was then performed to summarize the harmonic coefficients and to identify the axes that captured most of the shape variation among individuals. The first two principal components (PCs), Shape-PC1 and Shape-PC2, which explained 83% of shell outline variation (Supplementary Material Fig. S2), were used as new shape descriptors in subsequent modeling.

### Shell structure and composition

Shell sections were cut longitudinally along the axis of maximum shell growth from both left and right valves with a water-cooled diamond saw (Gryphon Aquasaw Bandsaw) and set in polyester resin (Supplementary Material Fig. S3). The embedded specimens were then sectioned with a high-speed diamond saw (Allied High Tech, TechCut 4), progressively ground (70−6 μm) and diamond polished (6−1 μm) using an automated grinder/polisher (Allied High Tech, M-Prep 5). Polished sections were imaged using a digital microscope (VHX-5000, Keyence) at LDEO. Images were processed and segmented with Fiji (v1.54b) and ilastik (v 1.4.0rc8) software (Berg *et al*., 2019) to measure the cross-section length and the areas of the whole section, foliated and chalk microstructures (Supplementary Material Fig. S3). Estimated folia and chalk areas were divided by the cross-section length to control for variability in shells’ profile and to calculate folia and chalk “thickness”. The proportion of chalk microstructure (chalk %) was measured as the ratio between chalk area and whole section area.

### Flexural properties

The flexural properties of the oyster shells were measured following the protocol developed by Wan, Ma & Gorb (2019). Rectangular beams (3 mm wide × minimal length of 26 mm) were cut from each of the left and right valves of fresh shells (Supplementary Material Fig. 4S), inspected to ensure beam integrity, wrapped in wetted degreasing cotton, and stored in plastic centrifuge tubes until testing. A Universal Testing Machine (Instron 5966, Instron, US) equipped with a fully articulated three-point bending fixture was used to break the beams and record measurements. Beams were mounted at the middle of their outer surface and load was applied at a strain rate of 0.001s^-1^ until fracture. A load-displacement curve was recorded for each sample. Following testing, fractographic examination was performed using a stereomicroscope (VHX-5000, Keyence) to investigate fracture patterns and measure the beam’s thickness at the fracture point. We used this information to calculate the Young’s Modulus (elastic modulus) (Supplementary Material Fig. S4) in accordance to the ASTM C1341-13 (ASTM, 2018) as in O’Toole-Howes *et al*., (2019).

### Shell density and bulk organics

After three-point bending tests, shell beams were used to calculate shell density and bulk organic abundance. We measured density using a fully automatic gas displacement (helium) pycnometer (AccuPyc 1330, Micrometrics). For bulk organic estimations, beams were first weighed and then ashed in ceramic crucibles for 5 h at 450°C using a high temperature furnace to burn organics. The ashed samples were then weighed again to determine the difference between initial shell mass and ash mass, which represents the bulk organics mass (organics %) (Telesca *et al*., 2019).

### Organic matrix content

Thermal gravimetric analyses (TGA) were performed to estimate the weight proportion (wt%) of the organic matrix (OM) within foliated and chalk microstructures, following the approach of Telesca *et al*. (2019). After removing the periostracum by sanding, fragments of individual microstructures were isolated along the posteroventral shell margin, air-dried, and finely ground. We tested 10 mg of these powdered samples with a thermogravimetric analyzer (TGA Q500, TA Instruments). The samples were heated at a rate of 10°C/min from 25°C to 600°C under a dynamic nitrogen atmosphere. Weight changes during heating were recorded with a precision microbalance. Based on literature values (Zaremba *et al*., 1998; Harper, Checa & Rodríguez-Navarro, 2009), as well as the known decomposition temperature of polysaccharides and proteins, three intervals of OM loss were identified and attributed to: *i*) the degradation of non-proteinaceous (polysaccharides) component of the inter-crystalline organic matrix (OM-I; 150-250°C), *ii*) the degradation of the proteinaceous inter-crystalline organic matrix (OM-II; folia: 250-430°C; chalk: 250-380°C), *iii*) the degradation of the intra-crystalline organic matrix (OM-III; folia: 430-550°C; chalk: 380-550°C) (Supplementary Material Fig. S5, Supplementary Methods S2).

### Crystal size, variability, and density

Shell material for scanning electron microscopy (SEM) was prepared to allow the examination of the internal surfaces of the valve. Shell samples were cut with a rotary tool from the posterior growing margin of each shell, bleached (10s in 1% NaOCl) to remove organic residues, rinsed, air-dried, mounted on aluminum stubs, and gold-coated (Cressington 108 Sputter Coater). Internal shell surfaces were imaged using a field emission SEM (FESEM; NanoSEM 450, FEI Nova; Supplementary Material Fig. S6). Calcitic crystals (laths) size, variability (standard deviation - SD), and density (number of crystals/unit area) were measured as in Meng *et al*. (2018).

### Magnesium content

The magnesium to calcium ratio (Mg/Ca) was analyzed on a ThermoScientific 6500 Inductively Coupled Plasma-Optical Emission Spectrophotometer (iCAP6500 ICP-OES, ThermoScientific) based on the technique utilized by Brenner *et al*. (2017). Powered samples of shell folia (5 mg) were subjected to wet chemical cleaning involving oxidation in hot (70°C) buffered H_2_O_2_–NaOH (0.1N NaOH, 15% v/v H_2_O_2_) solution to remove residual organic matter. Shell powder aliquots (∼300 μg) were diluted to 50 ppm Ca in 2% trace metal grade HNO_3_ and vortexed until no solid was visible in each vial. Solution standards were measured bracketing samples (every-other) to correct for instrumental drift and matrix effects resulting from variability in Ca concentration (Schrag, 1999). All values were drift corrected based on our internal standard, a diluted Matrix Match (MaMa) solution based on average oyster composition, with Mg/Ca at 5 mmol/mmol.

### Crystal elasticity and hardness

To measure microscale mechanical properties, we used a nanoindentation approach following the protocols of Karimzadeh *et al*. (2019) and O’Toole-Howes *et al*. (2019). Nanoindentation tests were performed on polished shell sections with a TriboIndenter Nanoindenter (Hysitron TI 950, Bruker) equipped with a diamond Berkovich tip. Well-polished regions of foliated structure were chosen using a light microscope on the nanoindenter, and a Scanning Probe Microscopy (SPM) map (60 × 60 µm) of the area was acquired before indentation. A 4×5 array with 10 µm spacing between indents was performed under displacement-controlled conditions at a loading/unloading rate of 10nm/s and a 20s hold at the maximum depth of 400 nm. Previous research showed that indentation depth has a significant effect on test results that are almost stable after 200 µm (Karimzadeh *et al*., 2019; O’Toole-Howes *et al*., 2019). A load-unloading curve was recorded for each sample and the Oliver-Pharr method (Oliver & Pharr, 2004) was used to determine hardness and reduced modulus. A Poisson’s ratio value of ν = 0.3 (O’Toole-Howes *et al*., 2019) was chosen to calculate the effective elastic modulus (Supplementary Material Fig. S7).

### PCA analysis

We employed a principal component analysis (PCA) approach to quantitatively evaluate the environmental and shell biomineralization variability (regimes) across the HRE, following the methodology outlined by Telesca *et al*. (2021). To obtain a first order estimate of the water conditions experienced by the sampled specimens throughout their lifespan, we calculated average growth season (i.e., June-October in the HRE) values for each parameter over a three-year period (2018-2020). We performed separate PCAs on *i*) the water quality parameters (i.e., temperature, salinity, DO, pH, Chl-a, turbidity), *ii*) macroscale shell traits (i.e., Shape-PC1, Shape-PC2, organics %, density, layer thickness, chalk %, elastic modulus) and *iii*) microscale shell traits (i.e., wt% organic matrix, elasticity modulus, hardness, Mg/Ca, crystal size, SD and density), to characterize environmental and biomineralization “regimes”. This approach was used to reduce the dimensionality of our datasets and to create a new set of independent summary variables, the principal components (PCs), capturing variability in water conditions and shell traits between locations. The first two PCs from each PCA were used as input variables to examine the effects of local environmental regimes on oyster macro and micro scale shell traits.

### Statistical analysis

Generalized linear models (GLMs) were used to compare macro and micro scale shell trait variations with respect to location and environmental regime. After data exploration (Zuur, Ieno & Elphick, 2010), variance inflation factors (VIFs) were calculated to check for collinearity between input variables. VIF values < 2 indicate an acceptable degree of correlation among covariates to be included within the same model. To directly compare model estimates (effect size metrics) from predictors on different measurement scales, we standardized (z-transformed) all the input variables (Schielzeth, 2010). Shell area was included in all models to control for possible effects of shell size (age) on the measured shell traits. Preliminary inspection of model residuals showed heteroscedasticity in most models. When the use of different continuous probability distributions (i.e., Gamma and inverse Gaussian) and link functions did not stabilize the variance, a ln-transformation of the response was required. Response variables did not require further transformations. To model proportional shell trait data (0 < *y* < 1), a GLM with a Beta distribution and a logistic link function was used. Models optimization and comparisons were performed using corrected Akaike Information Criterion (AICc) and bootstrapped likelihood ratio tests. The fixed component was optimized by rejecting only non-significant interaction terms that minimized the AICc value. The proportion of variance explained by the models was quantified with pseudo determination coefficients, pseudo-R^2^ (Nakagawa, Johnson & Schielzeth, 2017). Final models were validated by inspection of residual patterns to verify GLM assumptions of normality, homogeneity, and independence. Optimal models were used to estimate the mean effect sizes of predictors on the response variable (Schielzeth, 2010). Ninety-five per cent confidence intervals (95% CIs) for the regression parameters were generated using bias-corrected parametric bootstrap methods (10,000 iterations) and used for statistical inference; if the CIs did not include zero, then the effect was considered significant. All data exploration and statistical modeling were performed in R (v4.2.2) (R Core Team, 2022).

## RESULTS

### Macro scale shell patterns

The EFA revealed significant shape differences among sampling locations (MANOVA: N = 222, Wilk’s λ = 0.59, approximate-F_3,200_ = 4.36, *P* < .0001). Oysters from sites MCB and LCL shells were generally rounder and wider compared to PBP and BTP oysters (Supplementary Material Fig. S8, Table S2).

Section analyses indicated significant variations in shell total thickness (F_3,142_ = 7.33, *P* = 0.0002) and weight (F_3,142_ = 7.66, *P* < .0001), among collection sites for shells of a given size (Fig. 2A; Supplementary Material Fig. S9, Table S3). Shell of oysters from relatively high salinity environments (LCL, PCP, BTP) were on average 40% thicker than those from low salinity environments (MCB). We identified significant differences in the thickness of foliated and chalky calcite among locations (folia: F_3,74_ = 31.45, *P* <.0001; chalk: F_3,74_= 5.12, *P* = 0.0023; Figs. 2A; Supplementary Material Fig S9, Table S3). In both valves, foliated calcite constituted the predominant structure (left: mean[SD] = 70.2%[13.8]; right: mean[SD] = 77.6%[11.3]; Fig. 2B; Supplementary Material Fig. S9). During shell thickening, folia were deposited 52% faster than chalk (folia: mean[SE] = 0.68 mm[0.05], t_148_ = 13.19, *P* < .0001; chalk: mean[SE] = 0.32 mm[0.05], t_148_ = 6.30, *P* < .0001; Fig.2B), leading to a progressive decrease of the chalk % with increasing total thickness (Figs. 2A-B). Shell size (area) was positively correlated with thickness in all layers (Supplementary Material Table S3), indicating thickening during growth.

**Figure 2.**
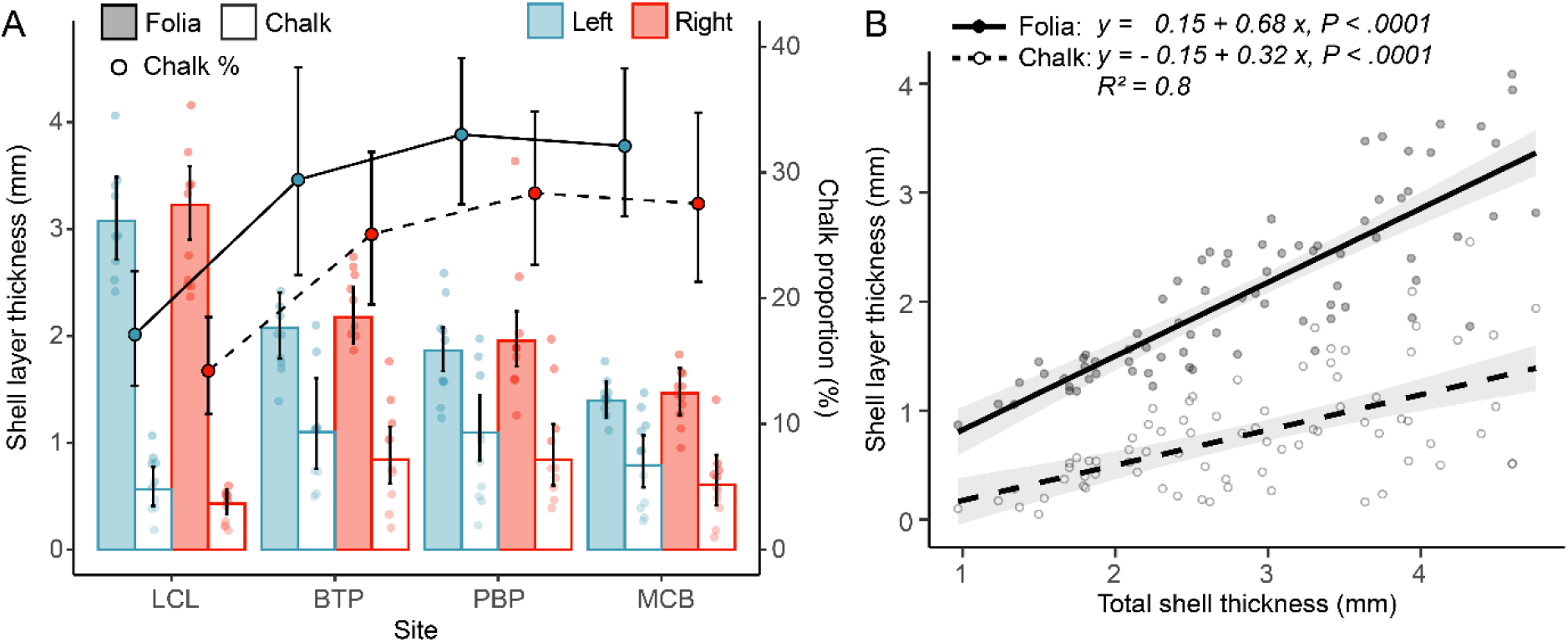
Spatial variations in oyster shell thickness and structure. **A**. Variation in the deposition of foliated (solid bars) and chalky (open bars) calcite in the left (blue bars) and right (red bars) valves at different sampling sites. We identified a general decrease in chalk % (solid circles) with increasing shell thickness in both valves. **B**. Relationship between the thickness of foliated (solid line) and chalky (dashed line) layers and the total shell thickness, showing significant differences in deposition rates (Total thickness × Layer: F_1,148_ = 23.74, *P* < .0001). The predicted values and their 95% CIs (error bars and shaded area) were estimated for oysters of a mean shell size (47 cm^2^).

We identified significant differences in elastic modulus (F_3,116_ = 4.71, *P* = 0.0039), shell density (F_3,112_ = 8.84, *P* < .0001) and bulk organics % (F_3,114_ = 15.35, *P* = 0.0015) between collection sites (Supplementary Material Fig. S10; Table S4). Low salinity oysters (MCB) displayed significantly higher elastic modulus, indicating formation of more brittle shells, compared to higher salinity oysters (Supplementary Material Fig. S10). We also observed a significant positive relationship between the elastic modulus and shell density, which differed between shell valves (Density × Valve: F_1,115_ = 5.52, *P* = 0.02; Fig. 3A), but no relationship with shell organics % (t_115_ = –0.88, *P* = 0.4; Supplementary Material Table S5). This indicates a stronger effect of inorganic rather than organic components on the elastic properties of oyster shells. Furthermore, density significantly decreased with increasing chalk % (z_117_ = –4.85, *P* < .0001; Fig. 3A; Supplementary Material Table S5).

**Figure 3.**
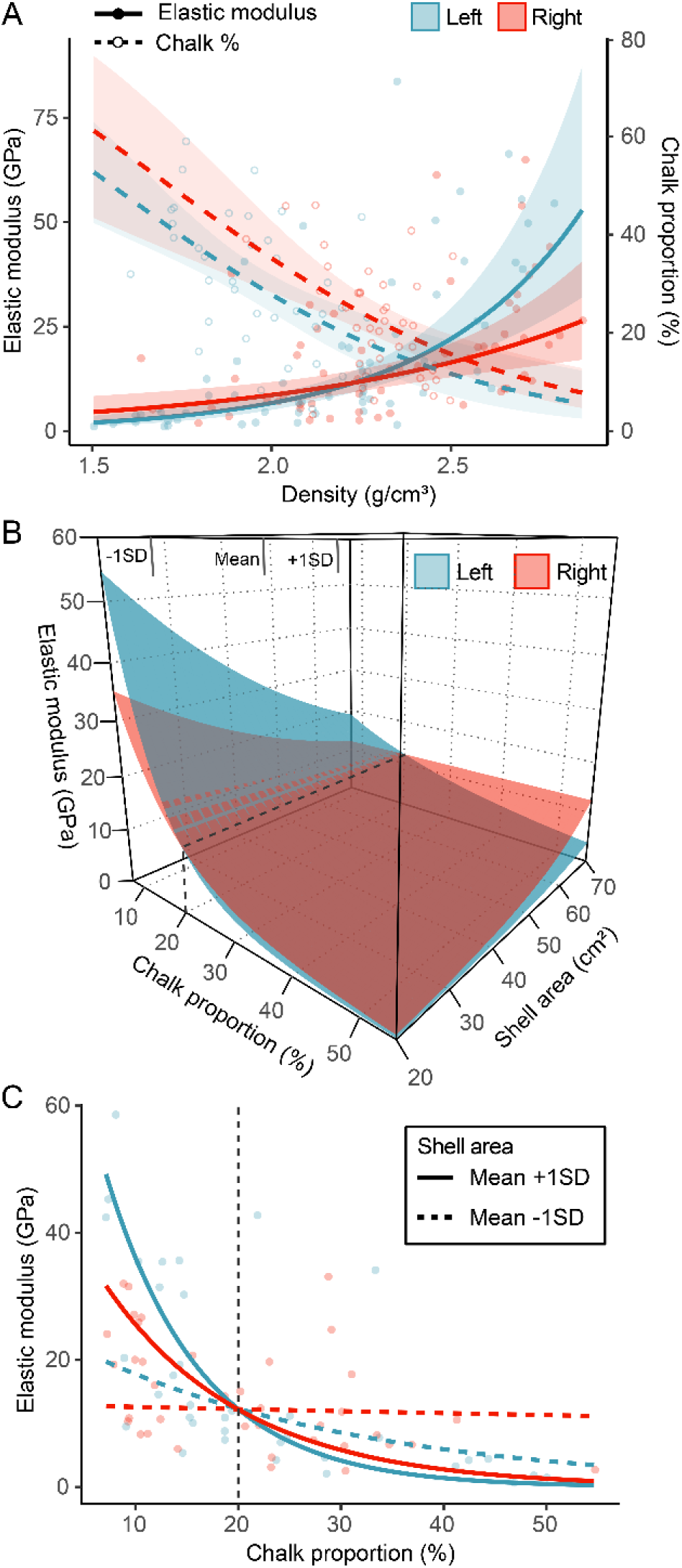
Predicted relationships between elastic modulus, density, and chalk %. **A.** The positive relationship between elastic modulus (solid lines) and density varied between the left (blue) and right (red) valves (left: mean[SE] = 3.90 GPa[0.71], t_115_ = 5.46, *P* < .0001; right: mean[SE] = 2.81 GPa[0.56], t_115_ = 5.05, *P* < .0001) and decreased with increasing shell size (Density × Area: t_115_ = –3.14, *P* = 0.002; Supplementary Material Table S5). The negative correlation between density and chalk % (dashed lines) underscores the strong impact of shell structure on flexural properties. Mean values and their 95% CIs (shaded areas) are predicted while controlling for shell size (47 cm^2^). **B.** Predicted multiple relationships between the elastic modulus of left (blue plane) and right (red plane) valves, chalk %, shell size, and their interactions (as in Eq. 1-2). Vertical marks indicate mean value ± 1SD of shell area. **C.** Interacting effects of chalk % and shell size on elastic modulus. The elastic modulus is modelled as a function of chalk % for the Mean ± 1SD values of shell area (i.e., 47±12 cm^2^), see panel **B**. Predictions (lines) indicate an inversion of the relative valve’s elastic modulus for chalk > 20% (vertical dashed line) and how this becomes less sensitive to changes in chalk % during shell growth.

The elastic modulus exhibited a strong inverse correlation with chalk %, but the strength of this relationship varied significantly between shell valves (Chalk% × Valve: F_1,69_ = 5.87, *P* = 0.015) and with size (Chalk% × Area: F_1,69_ = 21.7, *P* < .0001), as expressed in Equation 1. This overall trend indicates production of less brittle shells as the chalk % increases (Fig. 3B, Table 1):

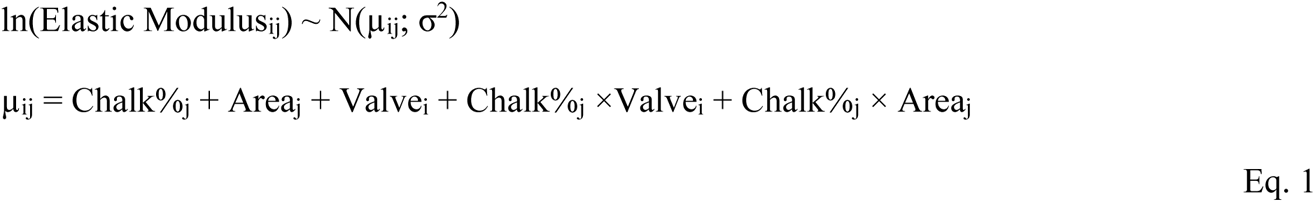

**Table 1.**
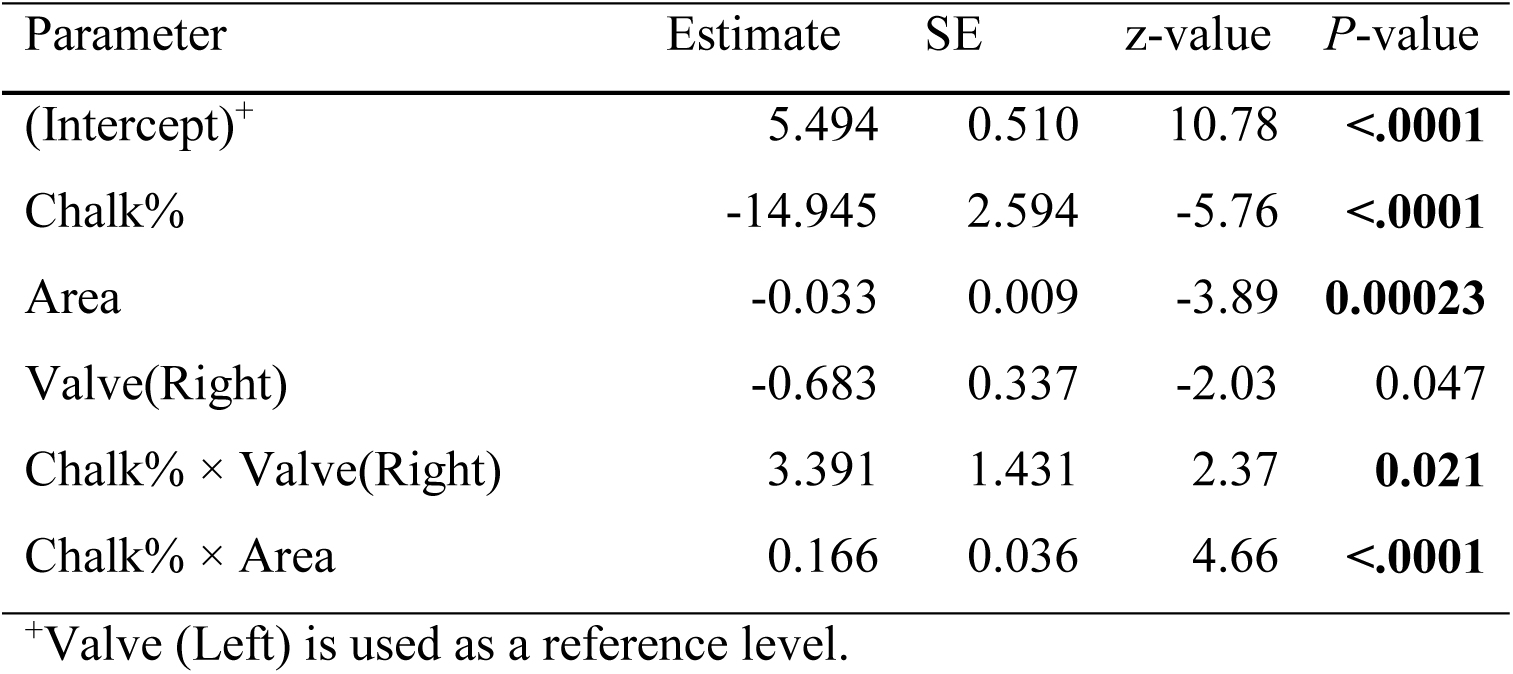
Elastic modulus GLM summary. Model summary statistics are reported for the modelled relationships between the elastic modulus, chalk % (*n* = 40) and shell size (Area), and how this changes between shell layers (folia, chalk) (Fig. 3B, C).

| Parameter | Estimate | SE | z-value | P-value |
| --- | --- | --- | --- | --- |
| (Intercept)<sup>+</sup> | 5.494 | 0.510 | 10.78 | <.0001 |
| Chalk% | -14.945 | 2.594 | -5.76 | <.0001 |
| Area | -0.033 | 0.009 | -3.89 | **0.00023** |
| Valve(Right) | -0.683 | 0.337 | -2.03 | 0.047 |
| Chalk% × Valve(Right) | 3.391 | 1.431 | 2.37 | **0.021** |
| Chalk% × Area | 0.166 | 0.036 | 4.66 | <.0001 |

<sup>+</sup>Valve (Left) is used as a reference level.

where Elastic Modulus_i*j*_ is the *j*th observation for valve *i* (*i* = left, right) that is assumed to follow a normal distribution with expectation *µ_ij_* and variance *σ^2^*. The effect size of chalk % on the elastic modulus of the left valve was 29% larger than that of the right valve, but the strength of this effect decreased with increasing shell size (Table 1; Eq. 2). This suggests a decreasing impact of chalk % on the mechanical properties of oyster shells during growth from small to large adults (Fig. 3C).

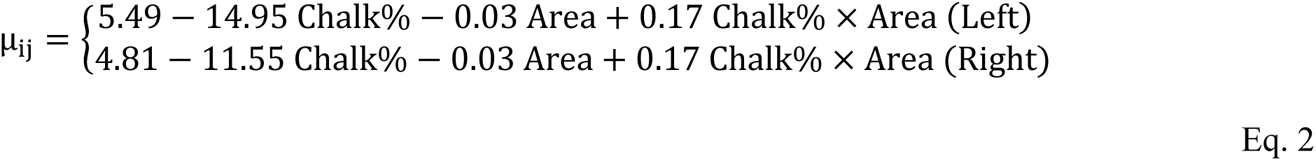

### Micro scale shell patterns

TGA results revealed that chalk was characterized by an average of 68% more weight proportion of organic matrix (OM wt%) compared to folia (chalk: mean[SE] = 3.2 wt%[0.1]; folia: mean[SE] = 1.9 wt%[0.1]), implying a lower calcification of chalk compared to folia (Fig. 4A). The OM wt% of folia significantly differed between collection sites, whereas no between-site variation in chalk % was detected (Fig. 4A; Supplementary Material Table S6). However, the wt% of inter-crystalline OM-I/II components significantly differed between locations in both folia and chalk (Fig. 4C; Supplementary Methods S2), indicating marked qualitative changes in OM components. The wt% of inter-crystalline OM-II showed an inverse relationship with intra-crystalline OM-III wt% (z_36_ = –8.08, *P* < .0001, pseudo-R^2^ = 0.85) in both layers (Supplementary Material Fig. S11).

**Figure 4.**
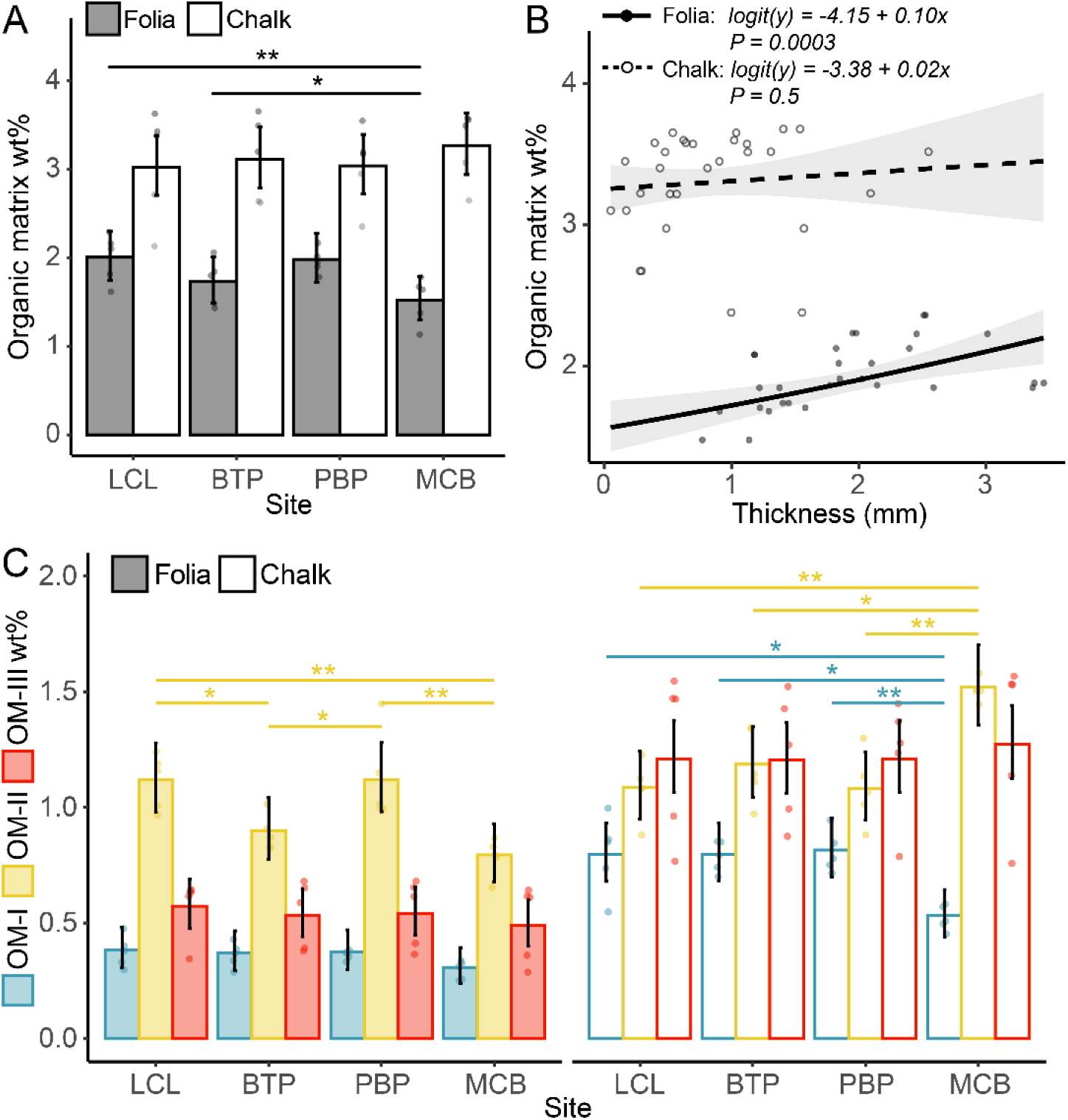
Shell OM wt% patterns. **A.** Variation of OM wt% between collection sites for folia (solid bars) and chalk (open bars). Within each pair, OM wt% is significantly higher in chalk than folia (mean[SE] = 1.3wt%[0.11], z_38_ = 12.11, *P* < .0001), in addition to significant difference variation between folia between sites. **B**. Predicted relationships between the wt% of OM and the thickness of foliated (solid line) and chalk (dashed line) layers. We identified a positive correlation of folia OM wt% with shell thickness (mean[SE] = 0.10[0.028], z_115_ = 3.66, *P* = 0.0003), but no trend for chalk (mean[SE] = 0.02[0.026], z_115_ = 0.68, *P* = 0.5), suggesting deposition of progressively less calcified folia (lower wt% of CaCO_3_) during layer thickening. **C.** Between site variations in organic matrix components: non-proteinaceous inter-crystalline OM-I (blue), proteinaceous inter-crystalline OM-II (yellow), proteinaceous intra-crystalline OM-III (red). Bar plot error bars and shaded areas represent 95% CIs. Significant pairwise contrasts and regression coefficients are reported as upper brackets (ns *P* >.5, * *P* < .05, ** *P* < .001, *** *P* < .0001; see Supplementary Material Table S6).

SEM imaging showed significant differences in mean size (χ^2^_3_ = 8.15, *P* = 0.04) and standard deviation (SD; χ^2^_3_ = 8.46, *P* = 0.03) of calcitic crystals (laths) between locations, whereas laths density (number of crystals/unit area) exhibited no significant variation (F_3, 16_ = 1.59, *P* = 0.2; Supplementary Material Fig. S12). No relationship emerged between crystal traits and shell size (size: t_16_ = −0.15, *P* = 0.8; SD: t_16_ = 0.04, *P* = 0.9; density: t_16_ = –0.94, *P* = 0.4; Supplementary Material Table S7). Crystal size was positively correlated to crystal SD (t_16_ = 8.91, *P* < .0001), but negatively correlated to crystal density (t_16_ = –4.92, *P* < .0001; Fig. 5A). We identified a significant positive relationship between crystal density and intra-crystalline OM wt% (z_114_ = 3.17, *P* = 0.002, pseudo-R^2^ = 0.81), while crystal size exhibited no correlation with OM wt% components (size: z_116_ = 1.28, *P* = 0.2, pseudo-R^2^ = 0.79). ICP analyses indicated marked differences in Mg/Ca ratio between sampling locations (F_3,33_ = 16.48, *P* < .0001). Specimens from the low-salinity MCB location displayed the highest Mg/Ca ratios (mean[SE] = 8.53 mmol/mol[0.35], in contrast to the average ratio observed in other locations (mean[SE] = 5.43 mmol/mol[0.19]; Supplementary Material Fig. S13, Table S8). Moreover, left valves exhibited an average Mg/Ca ratio 19% higher than right valves (mean[SE] = 1.09 mmol/mol[0.28], t_33_ = 3.95, *P* = 0.0004).

**Figure 5.**
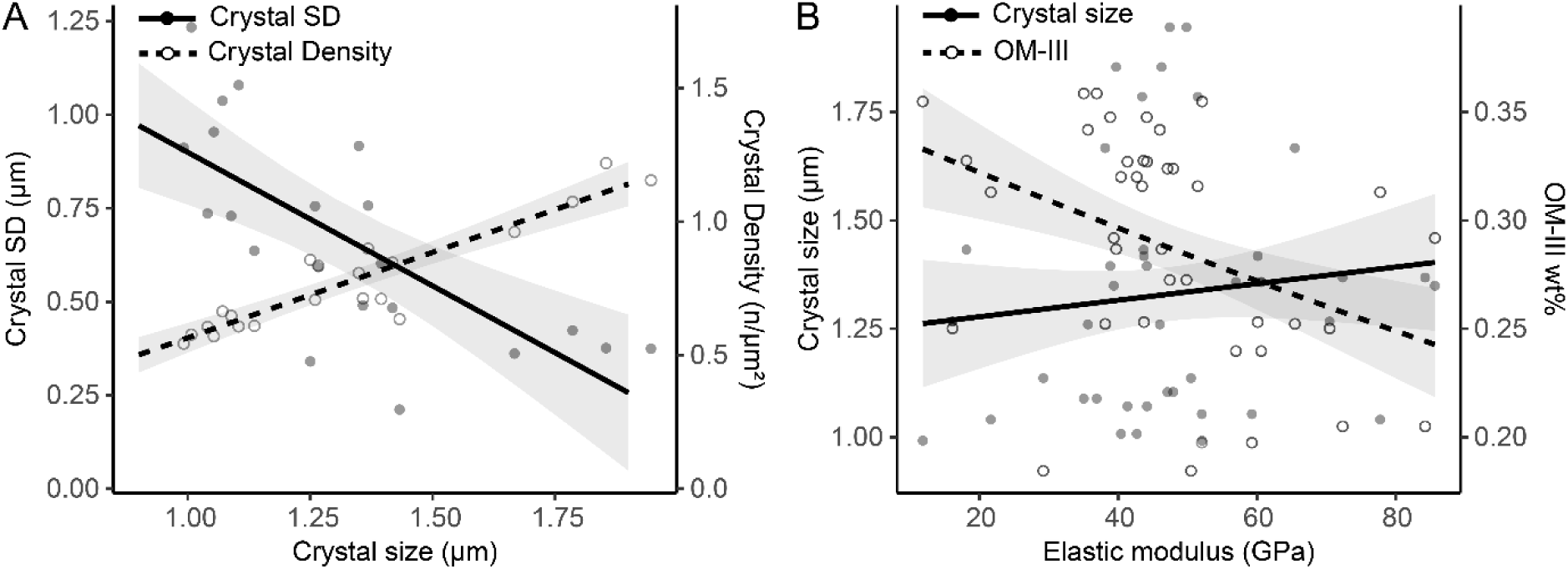
Predicted relationships between crystal traits, elastic modulus, density, and organic content. **A.** Significant covariations of crystal size with crystal SD (solid line) and crystal density (dashed line). **B.** Relationships of elastic modulus with crystal size (solid line; mean[SE] = 3.42 GPa[6.15], t_77_ = 0.55, *P* = 0.6) and OM-III (dashed line; mean[SE] = - 119.18 GPa[33.17], t_77_ = –3.59, *P* = 0.00057), indicating a more pronounced influence of OM content (both OM-III and OM-II wt%; see Supplementary Material Fig. S11) compared to calcitic crystal traits on the elastic modulus. The predicted values (line) and their 95% CIs (shaded areas) were estimated for oysters of a mean shell size (47 cm^2^).

Nanoindentation analyses revealed significant variation in elastic modulus (F_3,601_ = 6.33, *P* = 0.00031) and hardness (F_3,601_ = 20.39, *P* < .0001) of foliated calcite between the sampling locations, but no difference between shell valves (Supplementary Material Fig. S13, TableS8). Elastic modulus showed a positive relationship with hardness in both valves (t_602_ = 13.25, *P* < .0001, R^2^ = 0.53; Supplementary Material Fig. S13). These findings contrast with the trend observed for macroscale elastic modulus (Supplementary Material Fig. S10), suggesting a potential trade-off between macro- and micro-scale flexural properties. We also identified a significant negative relationship between the elastic modulus and shell OM-III wt%, but no relationship with crystal size was detected (Fig. 5B). Finally, we observed a marginally significant negative relationship between crystal hardness and Mg/Ca ratios (mean[SE] = –0.07 GPa[0.04], t_77_ = –2.02, *P* = 0.047; Supplementary Material Fig. S8, Table S13).

### Environmental and shell biomineralization regimes

The PCA on water quality parameters revealed variations in environmental regimes across the sampling locations (Supplementary Material Fig. 6A, Table S9). Envrio-PC1 (72.8%) captured variations in chemical and biological water parameters: turbidity, pH, DO and Chl-a concentration. Increasing Enviro-PC1 values corresponded to declining values of these variables. Enviro-PC2 (21.9%) reflected variations in physical water parameters, indicating a strong impact of salinity (ranging from 7 to 25 PSU) and a milder influence of temperature (ranging from 22 to 23 °C). Increasing Enviro-PC2 values indicated diminishing impacts of salinity but increasing temperature.

The PCA on macroscale shell traits (Fig. 6B; Supplementary Material Table S9) indicated that Macro-PC1 (29.38%) captured variations in shell elongation, total and foliated thickness, density, organic %, and elastic modulus, while Macro-PC2 (25.7%) captured variations in chalk thickness and chalk %. Increasing Macro-PC1 values indicate deposition of thicker (both total and foliated thickness), rounder, and more ductile shells, while increasing Macro-PC2 values corresponded to decreasing thickness and relative abundance of chalky calcite.

**Figure 6.**
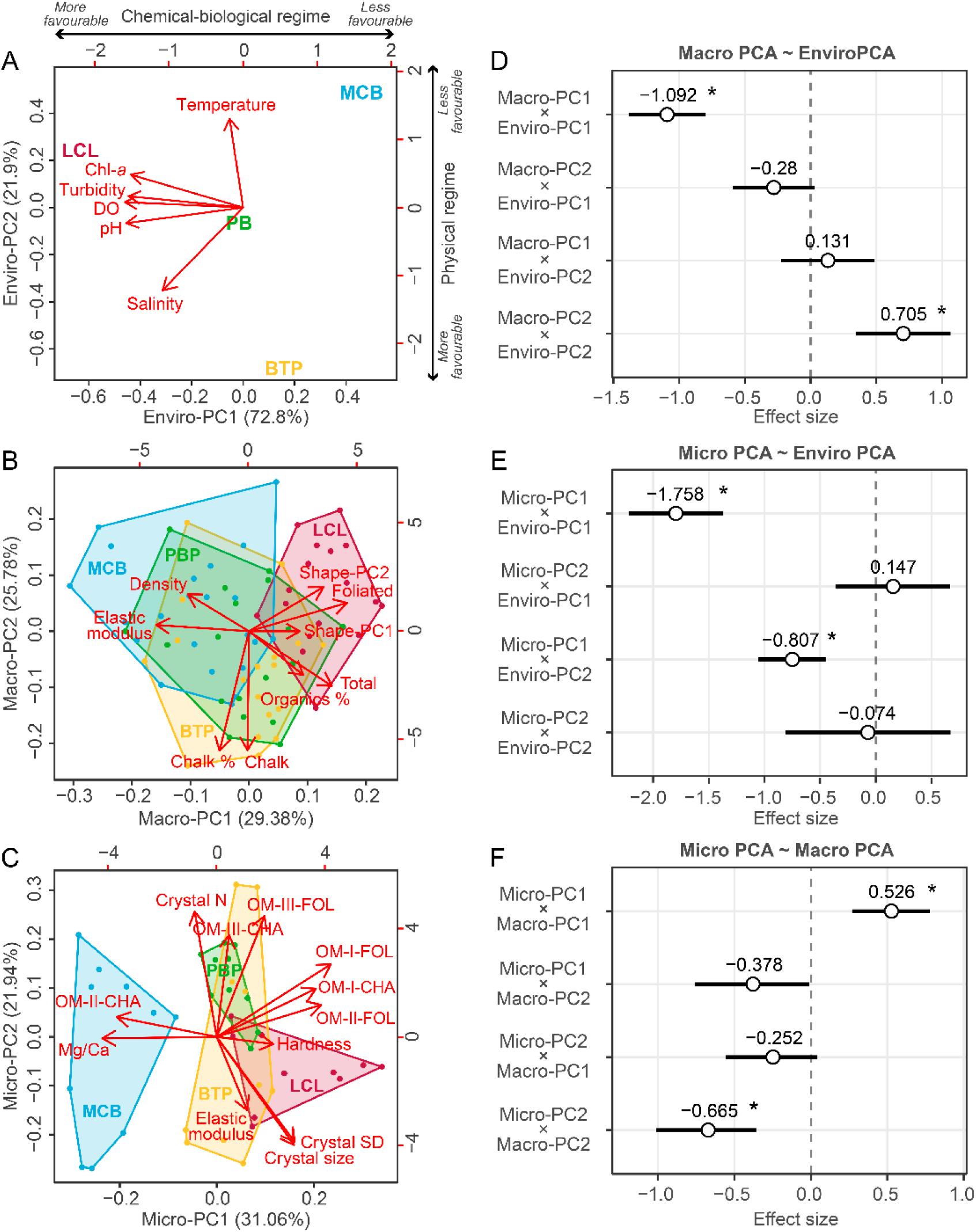
PCA biplot of **A.** environmental regimes, **B.** macro scale and **C.** micro scale shell trait variations between study locations. Top and right axes report the loading scores (i.e., the coefficients of the linear combination of the original variables used to construct the PC). Red arrows represent the loading vectors of the original variables on the PCs, showing how each descriptor is correlated with the plotted PCs (Supplementary Material Table S9), and point the direction of increasing value for the original variables. The placement of locations on the plane is determined by their relative PC scores and indicates differences in environmental and biomineralization regimes. (Total = size-corrected total shell thickness, Chalk = size-corrected chalk thickness, Foliated = size-corrected folia thickness, Crystal N = crystal number/unit area, Crystal SD = crystal size standard deviation). **D-F.** Mean effect size for coefficients estimated from pairwise GLMs between shell trait PCs and environmental PCs. Effect sizes (points), bootstrapped 95% CIs (error bars), and direction of change (coefficient values) are estimated for individual linear models (Table 1): **D.** Macro-PCs ∼ Enviro-PCs, **E.** Micro-PCs ∼ Enviro-PCs, and **F.** Micro-PCs ∼ Macro-PCs. Significance of regression parameters is identified when the 95% CI does not cross zero (* denotes significance).

The PCA on microscale shell traits (Fig. 6C; Supplementary Material Table S9) indicated that Micro-PC1 (31.06%) captured variations in shell hardness, Mg/Ca ratio, and inter-crystalline OM-I/II wt%, while Micro-PC2 (21.94%) captured variations in elastic modulus, intra-crystalline OMIII wt%, and crystal traits. Increasing Micro-PC1 values corresponded to formation of harder microstructures, enriched in OM-I/II and with lower Mg/Ca ratios, while increasing Micro-PC2 values correspond to the formation of less brittle shells with an increased deposition of OM-III, and smaller calcitic crystals that are denser and more homogeneous.

### Summary of environmental control on shell traits

The observed significant relationships between Macro-PC1 and Enviro-PC1 and between Macro-PC2 and Enviro-PC2 indicated significant responses of macroscale shell traits to variations of physical, chemical, and biological water parameters (Fig. 6D; Table 2). Micro-PC1 showed a significant relationship with both Enviro-PC1 and Enviro-PC2 (Fig. 6E; Table 2), but the effect size of Enviro-PC1 (mean[SE] = –1.76[0.16]) was about 2.4 times larger than that of Enviro-PC2 (mean[SE] = –0.81[0.10]). No relationship between Micro-PC2 and environmental regimes was detected (Fig. 6E; Table 2).

**Table 2.**
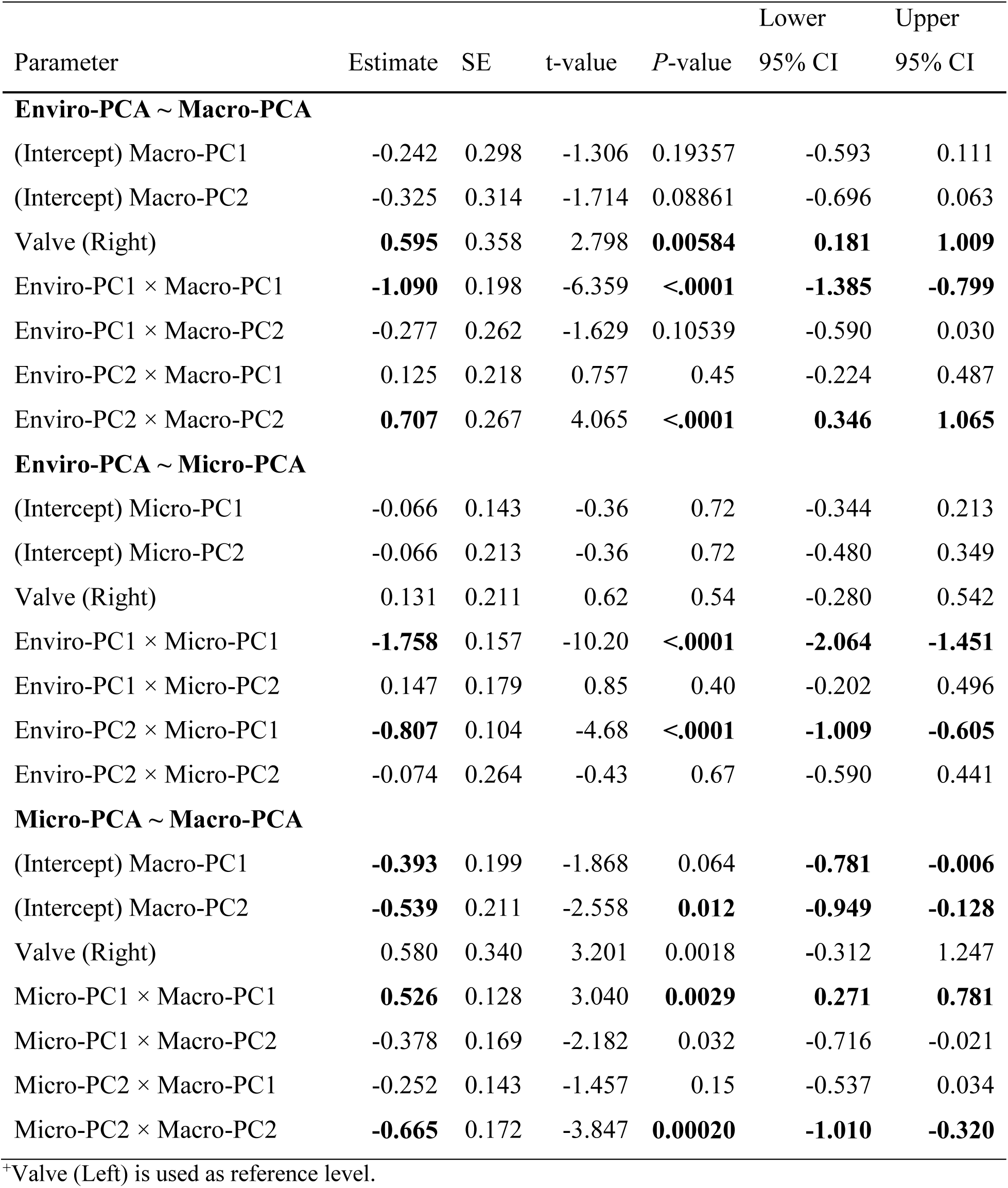
Summary of Enviro-, Macro- and Micro-PCs models. GLMs summary statistics and bootstrapped 95% confidence intervals (CIs) for the modelled relationships between Enviro-PCs, Macro-PCs and Micro-PCs (Fig. 6D-F) and among shell valves (Valve). Estimates are reported separately for each PC. Parameters’ significance (bold values) is determined when the 95% CI does not include zero.

The correlation between Macro-PC1 and Micro-PC1 was strongly positive (Fig. 6F; Table 2), which aligns with their co-variation with Enviro-PC1. Conversely, Macro-PC2 showed a strong negative relationship with Micro-PC2 (Fig. 6F; Table 2). The lack of covariation between Micro-PC2 and environmental regimes suggests that Macro/Micro-PC2 correlation indicates compensatory responses of shell microstructural traits to macrostructural alterations.

## DISCUSSION

Our results provide novel insights into how a reef-building oyster maintains shell integrity and function along natural environmental gradients by adaptively adjusting shell production, structure, and composition at the macro and micro scales. An understanding of the biological mechanisms and drivers of variation in natural environments is crucial to accurately predict population responses in highly variable and rapidly changing ecosystems (Urban *et al*., 2016). Estuarine restoration programs offer unique opportunities by providing a natural laboratory to investigate population-level responses under naturally complex field conditions and standardized growth conditions (i.e., restoration timing, substrate used, depth, and genetic pool). In contrast to laboratory studies, these natural organismal responses are a function of estuarine ecological complexity (Kroeker, Kordas & Harley, 2017) and incorporate transgenerational phenotypic plasticity (Cross, Harper & Peck, 2018; Telesca *et al*., 2021), both key to longer term adaptation and resistance.

### Estuarine patterns of oyster shell production

*Crassostrea virginica* shows marked changes in shell structure, and protective capacity (mechanical performance) with changing thickness under different salinity regime (Figs 2-3B). Specifically, as shell thickness increases with salinity, we observe a progressive shift in shell structure from production of thin, brittle shells with a higher proportion of foliated calcite at low salinity, to production of thicker, mechanically stronger shells with an increased proportion of chalky calcite under high salinity (Fig. 2). Despite these structural shifts, folia are always relatively more abundant than chalk during shell growth (Fig. 2B).

The observed decrease in shell production with salinity is in line with documented patterns of bivalve calcification under changing CaCO_3_ chemistry (Thomsen *et al*., 2018; Telesca *et al*., 2019). Specifically, the stability and deposition rate of bivalve shells depend on both the CaCO_3_ saturation state of seawater (i.e., Ω = [Ca^2+^][CO ^2−^]/*K*sp; with *K*sp = stoichiometric solubility product (Mucci, 1983)) and the ratio of calcification substrates ([Ca^2+^] and [HCO_3_ ^−^]) to the inhibitor (H^+^), expressed as the extended substrate–inhibitor ratio (ESIR = [Ca^2+^][HCO_3_^−^]/[H^+^]) (Sanders *et al*., 2021). The concentration of seawater HCO_3_^−^ and Ca^2+^ decreases with salinity, leading to lower Ω of CaCO_3_ and ESIR (Thomsen, *et al*., 2015; Sanders *et al*., 2021). Low Ω_CaCO3_ constrains shell formation (Waldbusser *et al*., 2015) and favors CaCO_3_ dissolution (Thomsen, *et al*., 2015; Ries *et al*., 2016), while low ESIR predicts reduced calcification rates (Thomsen *et al*., 2018; Sanders *et al*., 2021). Our results suggest that at low salinities oyster calcification rates are likely impeded by low Ω and ESIR, resulting in the formation of thinner shells under more corrosive conditions. Conversely, saltier waters are less likely to constrain CaCO_3_ stability and calcification, favoring the formation of thicker oyster shells.

Observed variations in the predominant shell structure produced could be explained by changes in the energetic cost of calcification with salinity (Sanders *et al*., 2018). At low salinities (<10 PSU), when [Ca^2+^] is increasingly limited, the higher costs of epithelial ion transport (McConnaughey & Whelan, 1997), coupled with kinetic constraints on mineral precipitation at the calcification site (Waldbusser, Hales & Haley, 2016), have been demonstrated to substantially increase the energetic cost of calcification (Sanders *et al*., 2018). Previous studies have determined that the proportion of the total assimilated energy used for shell growth (both CaCO_3_ and organic matrix production) increased from less than 10% under fully marine conditions (Watson, Morley & Peck, 2017) to 36% to 67% at salinities ranging 6-16 PSU (Sanders *et al*., 2018). Although bivalve shells contain an average of ∼5% OM, its formation is significantly more costly (29 J mg^−1^) than that of CaCO_3_ production (average ∼1.5 J mg^−1^) (A. R. Palmer, 1992; Waldbusser *et al*., 2013). Following this, our observations suggest that the higher OM content of chalk (Fig. 4A) makes it energetically more expensive to produce than folia. However, the OM content and, therefore, the net energetic cost of folia approximately doubles with increasing shell thickness (Fig. 4B). Therefore, we suggest that to sustain shell growth under low salinity conditions, where both calcification and shell production costs are high, *C. virginica* favors production of thin shells predominantly made of energetically cheaper (i.e., low OM wt%) foliated calcite. Conversely, decreasing calcification costs at higher salinity favors production of thicker shells with an increasing proportion of energetically expensive chalk but a decreasing proportion of OM-enriched folia.

Our results further demonstrate that changes in oyster shell structure from small to large adults has a strong but variable impact on the shell’s mechanical performance (Fig. 3B-C), which is in line with estuarine gradients of predation pressure and size-dependent vulnerability of oysters to predation. Salinity positively correlates with abundance and predation pressure of shell crushing (durophagous; e.g., crabs, fish) and drilling (e.g., whelks) oyster predators (Zachary & Haven, 1973; Kimbro *et al*., 2017; Pusack *et al*., 2019). Although some drilling predators selectively consume large oysters (50-100 mm) (Pusack *et al*., 2018), oyster susceptibility to predation generally decreases during shell growth (Juanes, 1992), with the highest predation frequencies on oysters with shell heights up to 45-50 mm (Bisker & Castagna, 1987; Roger I E Newell, Kennedy & Shaw, 2007). As a response to predation, eastern oysters have shown a strong capacity for phenotypic plasticity in shell traits (Lord & Whitlatch, 2012; Robinson *et al*., 2014; Scherer & Smee, 2017). Previous studies have demonstrated different mechanical behaviors of oyster microstructures (S. W. Lee, Kim & Choi, 2008; S.-W. Lee *et al*., 2011). For instance, folia are characterized by high hardness and stress absorption (S. W. Lee, Kim & Choi, 2008), while the lower density of chalk makes it less hard but capable of greater crack/force absorption than folia (S.-W. Lee *et al*., 2011). In the HRE oysters, we reveal a strong inverse relationship between shell elastic modulus and both chalk abundance and shell size (Fig. 3B). In low salinity conditions (i.e., low predation), oysters produced thin shells with a low proportion of chalk, corresponding to a higher elastic modulus. This indicates shells that are brittle (i.e., less able to absorb energy) and with poor toughness (i.e., low resistance to crack propagation). On the other hand, oysters produced more durable, thicker shells with a higher proportion of chalk and lower elastic modulus at higher salinity (i.e., higher predation pressure) (Fig. 3B). However, the strength of the observed relationship between elastic modulus and chalk content decreases with increasing shell size (Fig. 3B-C). Indeed, we found that structural changes had a much stronger effect on the mechanical performance of small oysters compared to larger ones, suggesting diminishing needs for physical protection once a refuge size (> 50 mm) (Bisker & Castagna, 1987) is attained. Furthermore, shells with rounder profiles provide increased compression resistance (Johnson, 2020) and are preferentially produced under higher salinity than the more elongated, weaker shells prevalent in low salinity environments (Supplementary Material Fig. S8). These patterns suggest a capacity of *C. virginica* for trade-offs between shell composition and mechanical performance as compensatory responses to differential predation pressure during development.

### On the functional value of chalk

Although our observational study deals with the drivers of oyster shell production and composition, it allows us to speculate on the functional properties of chalk. Chalky calcite is a unique microstructure among molluscan shells that is found only in true oysters (Family Ostreidae). Although we have detailed descriptions of chalk’s morphology, crystallography, and formation (R. Palmer & Carriker, 1979; Higuera-Ruiz & Elorza, 2009; Checa, Harper & González-Segura, 2018), its exclusive presence in oyster shells has sparked a variety of opinions as to whether chalk formation is *i*) a way of maintaining a smooth interior shell profile while growing on irregular substrates (Korringa, 1951; Galtsoff, 1964; Checa, Harper & González-Segura, 2018), *ii*) a consequence of low or high CaCO_3_ saturation state (Margolis & Carver, 1974), *iii*) a result of the mantle detaching from the shell (Orton, Amirthalingam & Bull, 1927), or even, *iv*) a form of “remote mineralization” controlled by bacterial activity (Vermeij, 2014; Banker & Sumner, 2020). However, little is known about chalk deposition and functions under changing salinity.

Building on previous studies that highlighted chalk’s ductility (S.-W. Lee *et al*., 2011; Lombardi *et al*., 2013), extrapolating our results (Fig. 3B) suggests that a hypothetical oyster shell made entirely of either folia or chalk would be too brittle or too ductile, respectively. However, the addition of chalk deposits to a predominantly foliated structure greatly improves its mechanical performance (S.-W. Lee *et al*., 2011) regardless of shell size and thickness. We find that at less than ∼20% chalk content (Fig. 3.B-C) shell brittleness increases dramatically with only small decreases in chalk abundance. When chalk content is above ∼20%, shell mechanical performance improves but becomes less sensitive to chalk content as oyster size increases, suggesting a potential adaptive value in chalk plasticity. Moreover, the strength of the right shell valve, which is subject to the highest predation frequency (Carriker & Van Zandt, 1972), shows the strongest change in sensitivity with shell size. We suggest that, regardless of the adaptive or exaptive (Gould & Vrba, 1982) nature of the observed plastic responses, chalk incorporation and its advantage for a shell’s mechanical resilience against predation pressure should play key roles in maintaining shell integrity under selective environments.

### Patterns of shell microstructure and properties

This study reveals marked changes in *C. virginica* shell organic matrix (OM) production, crystal traits, and Mg/Ca (Figs 4-5), indicating a strong capacity of oysters to adjust their microstructure and, thereby, improve mechanical and dissolution resistance. While *C. virginica* shells are made of the chemically more resistant calcite polymorph of CaCO_3_ (Mucci, 1983), other factors, including the abundance of the OM, crystal size and orientation, as well as Mg/Ca ratios, have been shown to place additional benefits and constraints on shell physical protection and solubility (Harper, 2000; Chadwick *et al*., 2019; O’Toole-Howes *et al*., 2019; Leung *et al*., 2020). For instance, bivalve shells contain a low, but significant, proportion of organic material distributed between and within individual crystals (Watabe, 1988). This OM is composed of insoluble chitin and proteins along with soluble sulfated- and muco-polysaccharides (Wheeler, 1992). The quantity, distribution, and composition of the OM are important because they can affect shell solubility by either *i*) protecting the CaCO_3_ crystals from contact with corrosive seawater (Harper, 2000), *ii*) releasing acid residues through preferential OM degradation (Glover & Kidwell, 1993), or even *iii*) facilitating crystal disaggregation by loss of binding organics (Peck *et al*., 2015). Moreover, the OM composition should impact its protective capacity, given the higher resistance to dissolution and stability of chitin and proteins compared to polysaccharides.

TGA analyses reveal marked variations in OM distribution and composition between sites (Fig. 4A, C), including changes in the relative abundance of inter- vs intra-crystalline OM, and differences in the abundance of non-proteinaceous and proteinaceous components. These changes indicate deposition of a higher proportion of more proteinaceous inter-crystalline OM in low-salinity environments (MCB location), suggesting an improved protection of calcitic shell crystals and a higher resistance to corrosion under low Ω_CaCO3_ (Waldbusser *et al*., 2015). Moreover, we find that increased deposition of intra-crystalline OM has a positive effect on shell durability and fracture resistance (decreased elastic modulus and hardness; Fig. 4B), which is in line with previous studies demonstrating the role of OM in improving elastic properties and energy dissipation in skeletal biocomposites (Kamat *et al*., 2000; Côté, Darkins & Duffy, 2015; Leung *et al*., 2020). Overall, the strong inverse relationship between inter- and intra-crystalline OM content (Supplementary Material Fig. S11), along with their benefits for corrosion resistance and mechanical performance, could indicate trade-offs between distribution and composition of OM as a response to specific selective requirements for physical or chemical protection.

Crystal size, morphology and surface-area-to-volume ratios have also been shown to affect dissolution rates (Harper, 2000). Since individual crystals are not fully exposed to dissolution when they are enveloped by OM, dissolution preferentially proceeds along crystal edges and is related to the relative surface areas of crystals when they outcrop on the shell exterior (Harper, 2000). Therefore, microstructures with larger crystals (i.e., smaller surface area/volume) and with low crystal density (no. of crystals/unit area), overall having less crystal surface exposed to seawater, should be more resistant to dissolution (Chadwick *et al*., 2019). Our results indicate strong covariation of these crystal traits (Fig. 5A) with formation of folia with large crystals and low density under low salinity, suggesting an increased resistance to dissolution. We also show that intra-crystalline OM wt% has a stronger effect on elastic modulus of folia than crystal traits (i.e., size, density (Fig. 5B), corroborating the importance of the organic phase in improving shells’ elastic properties (Kamat *et al*., 2000).

An increasing proportion of magnesium (Mg^2+^) incorporated in the calcite lattice is known to generally increase both chemical solubility and hardness of calcite (Bischoff, Mackenzie & Bishop, 1987; Kunitake, Baker & Estroff, 2012). Surprisingly, we find increased Mg/Ca ratios under low salinity (MCB location) and a marginally significant but negative relationship between Mg/Ca ratio and shell hardness (Supplementary Material Fig. S13). This trend is puzzling because a higher Mg/Ca ratio in oysters would increase solubility (Bischoff, Mackenzie & Bishop, 1987), exacerbating dissolution in low-salinity waters with low Ω_CaCO3_. As previously suggested for high-magnesium coralline algae (Ragazzola *et al*., 2020), a higher substitution rate of Mg^2+^ for Ca^2+^ might be a consequence of a general lack of Ca^2+^ (Thomsen *et al*., 2018). In our oysters, this trend appears to be compensated by increased deposition of protein-enriched, inter-crystalline OM (Fig. 4C). Although previous studies have shown a positive relationship between Mg/Ca and hardness in calcite (Kunitake, Baker & Estroff, 2012), our interpretation is supported by molecular simulation studies, suggesting that amino acid content (i.e., OM proteins) in calcite can have a larger but opposite effect to Mg^2+^ on hardness and elasticity (Côté, Darkins & Duffy, 2015). Overall, compensatory increases of OM in our oysters may have overridden any Mg/Ca ratios effect on shell hardness.

### Chemical-biological and physical regime impacts on biomineralization and biomechanics

Our analysis of environmental regimes (Fig. 6A) allowed us to describe natural responses of *C. virginica* shell production, structure, and composition at the macro and micro scales (Figs 6C-B, 7) as a function of environmental controls (see also Kroeker *et al*., 2016; Telesca *et al*., 2021). We find that eastern oysters show a strong protective capacity for increased structural integrity and/or chemical resistance under variable calcification and predation regimes (Fig. 7).

**Figure 7.**
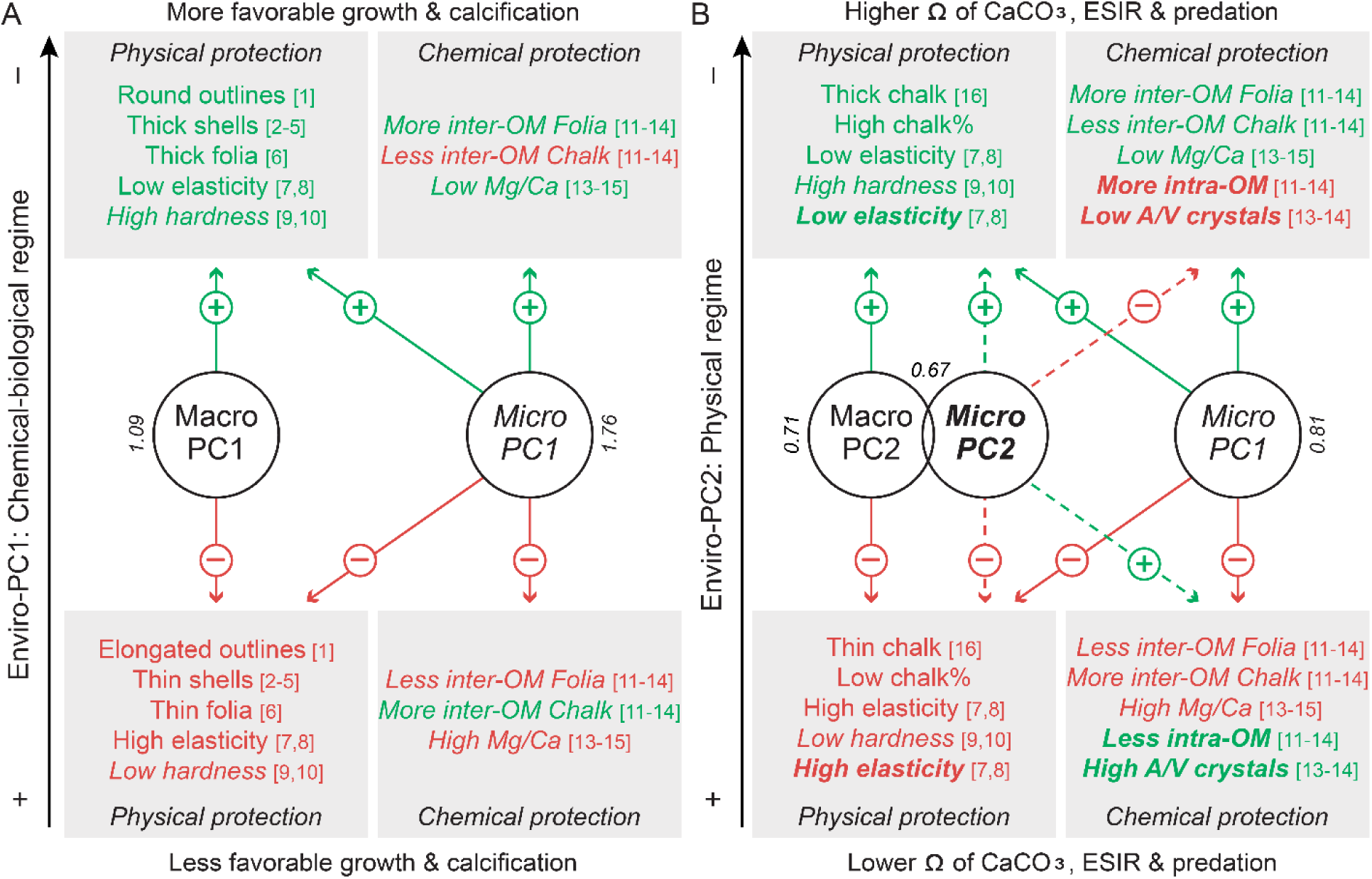
Conceptual representation of *C. virginica* quantitative and qualitative responses of macro and micro scale shell traits with changing environmental regimes in the HRE and their inferred contribution to physical and chemical protection provided by the shell. The graphs represent responses of macroscale (Macro-PC1/PC2; Fig. 6B) and microscale (Micro-PC1/PC2; Fig. 6C) shell traits to different **A.** chemical-biological (Enviro-PC1) and **B.** physical (Enviro-PC2) environmental regimes. Arrows indicate a positive (green; +) or negative (red; –) contribution to physical and chemical protective shell properties. Solid and dashed lines indicate direct and compensatory responses, respectively. Character type is used to differentiate contributions of Macro- and Micro-PCs (regular: Macro-PC1/2; italic: Micro-PC1; bold: Micro-PC2). Reported values represent effect sizes as in Figures 6D-F. Cited references supporting these effects are [1] Johnson (2020), [2] Gaylord *et al*. (2011), [3] Roger I. E. *et al*. (2007), [4] Large & Smee (2013), [5] Scherer *et al*. (2016), [6] S.-W. Lee *et al*. (2011), [7] Beniash *et al*. (2010), [8] Meng, Guo *et al*. (2018), [9] Meng, Fitzer *et al*. (2018), [10] Fitzer *et al*. (2015), [11] Kawaguchi & Watabe (1993), [12] Glover & Kidwell (1993), [13] Harper (2000), [14] Chadwick *et al*. (2019), [15] Bischoff, Mackenzie & Bishop (1987), [16] S.W. Lee, Kim & Choi (2008).

Our data suggest that chemical (pH and dissolved oxygen concentration) and biological (turbidity and Chl-a concentration, used as proxies for food supply) water conditions (Enviro-PC1) exert a significant effect on shell shape, deposition, and elastic modulus. These parameters also have a 179% greater effect than physical conditions on shell OM content, Mg/Ca ratios, and hardness (Fig. 6E). Increasingly favorable chemical-biological regimes (decreasing Enviro-PC1 values: increasing DO, food supply and pH), result in the formation of rounder and thicker shells, thicker folia, and decreasing elastic modulus (decreasing Macro-PC1), along with more abundant inter-crystalline OM, lower Mg/Ca ratios and increasing hardness (increasing Micro-PC1) (Fig. 7A). These observations suggest that oysters grow more physically and chemically durable shells under increasingly favorable regimes for oyster physiology and calcification.

Physical water conditions (Enviro-PC2: mostly salinity, with less significant temperature contributions) have a strong effect on chalk deposition, along with minor effects (compared to biological-chemical regimes) on changes in OM content, Mg/Ca ratios and hardness (Fig. 7B). With increasing salinity conditions (decreasing Enviro-PC2 values), oysters produce shells with relatively more calcitic chalk (decreasing Macro-PC2) and intra-crystalline OM (although the proportion of proteinaceous components in chalk OM decreases), decrease Mg incorporation, and increase hardness (increasing Micro-PC1). These observations suggest that the formation of more durable shells (i.e., low elastic modulus) and increased resistance to dissolution come at greater energetic expense. We also observe compensatory microstructural responses (increasing Micro-PC2) to increasing chalk production (decreasing Macro-PC2), showing formation of smaller, densely packed crystals, with more abundant intra-crystalline organic matrix (therefore decreasing OM-II; Supplementary Material Fig. S11), and lower elastic modulus. These compensatory adjustments contribute to increased mechanical performance (i.e., increased toughness, hardness, and intra-crystalline OM), but decrease resistance to dissolution (i.e., less proteinaceous intra-crystalline OM, and higher surface area/volume crystals) under marine conditions, while the reverse would be true in more brackish conditions.

In summary, *Crassostrea virginica* shows a strong capacity to respond to physical environmental demands via macrostructural changes in shell production and compensatory adjustments at the microstructural level (Fig. 7B). Under marine conditions, with higher Ω_CaCO3_, ESIR, and low dissolution levels, but high predation pressure, oysters produce shells that are physically stronger but chemically weaker. Conversely, in brackish waters, where chemical solubility is higher (i.e., low Ω and ESIR, along with limiting [Ca^2+^]), but predation pressure is significantly diminished, oysters produce shells providing increased chemical protection at the price of decreased structural integrity.

### Compensatory mechanisms in a changing environment

As hypothesized, our findings demonstrate a strong capacity of a restored population of *C. virginica* to withstand a wide range of environmental variability through unexpected levels of compensatory adjustments in shell biomineralization and biomechanical properties. These responses indicate salinity as a key predictor of eastern oyster shell formation. In marine conditions, where predation is high and calcification is favored, oysters grow thick shells with higher mechanical resistance but increased solubility, while under brackish conditions, where predation is reduced, or absent, and calcification is constrained, oysters produce more brittle shells with higher protection from dissolution. Increasing evidence supports the potential for resistance of calcifying organisms, including brachiopods (Cross, Harper & Peck, 2019), mytilid bivalves (Telesca *et al*., 2019), gastropods (Leung, Russell & Connell, 2017; Mayk *et al*., 2022), and coralline algae (McCoy & Ragazzola, 2014) to a range of biotic and biotic selective forces. This suggests compensatory biomineralization mechanisms (Telesca et al., 2021) to disturbance could represent a cornerstone for calcifiers to maintain calcifications and ecological functions under rapidly changing climate.

Our results also demonstrate the role of local-scale estuarine salinity gradients in shaping compensatory responses of biomineralization in adult oysters that we often assume to be less sensitive than early life-history stages according to experimental models (Waldbusser *et al*., 2015; Scherer *et al*., 2016). However, as the impacts of sea-level rise and climate change on estuaries accelerate and are exacerbated by growing anthropogenic influence on nearshore biogeochemical cycles (Levin *et al*., 2015; Röthig *et al*., 2023), rapid shits in estuarine salinity regimes may threaten this compensatory capacity of oysters. Molluscs shell calcification and mechanical properties face a range of emerging estuarine stressors, including acidification, elevated *p*CO2 levels, hypoxia, and limitation of food supply (Beniash *et al*., 2010; Melzner *et al*., 2011; Ivanina *et al*., 2013; Casas *et al*., 2018). While our finding demonstrate that *C. virginica* possesses great plasticity in shell calcification (Fig. 7), the increasing quantity and intensity of stressors, coupled with fluctuations in estuarine salinity (Röthig *et al*., 2023), pose a potential threat for *C. virginica* susceptibility in multiple stressors scenarios (Breitburg *et al*., 2015). In this regard, a better understanding of how natural processes shape or constrain compensatory responses of calcifying foundation species in naturally-complex environments, as the one presented in this work, is critical to anticipate changes to their functional roles in communities and, thereby, prevent damage to supported ecosystems and services in a rapidly changing world.

## ACKNOWLEDGEMENTS

We thank the Billion Oyster Project for making possible oyster collections at restored reefs in the NYC Harbor, and Dr. Jennifer Zhu (BOP) and Dr. Zofia Baumann (BOP - University of Connecticut) for support with samplings. We are grateful to Dindo Q. Mijares (NYU College of Dentistry) and Vasudev V. Nayak (University of Miami) for training on and support with flexural and nanoindentation testing. We thank the Columbia Nano Initiative for use of their Shared Lab Facilities, specifically thermogravimetry and SEM imaging. We also thank Prof. Reinhard Kozdon (LDEO) for the use of polishing laboratory equipment for sample preparation. The work was funded by the LDEO (Fellowship allowance), the LDEO Climate Center, and PADI Foundation under grant agreement no. PG011530.

## AUTHORS’ CONTRIBUTIONS

L.T., B.K.L., and B.H., conceived the original project; L.T. performed specimen collection, sample processing, laboratory work, data analysis and output; L.W. provided facilities and support for mechanical testing; L.T. prepared the first draft of the manuscript, and all co-authors contributed substantially to revisions.

