## Supplementary for "Biomineralization and biomechanical trade-offs under heterogeneous environments in the eastern oyster *Crassostrea virginica*"

SUPPLEMENTARY FIGURES

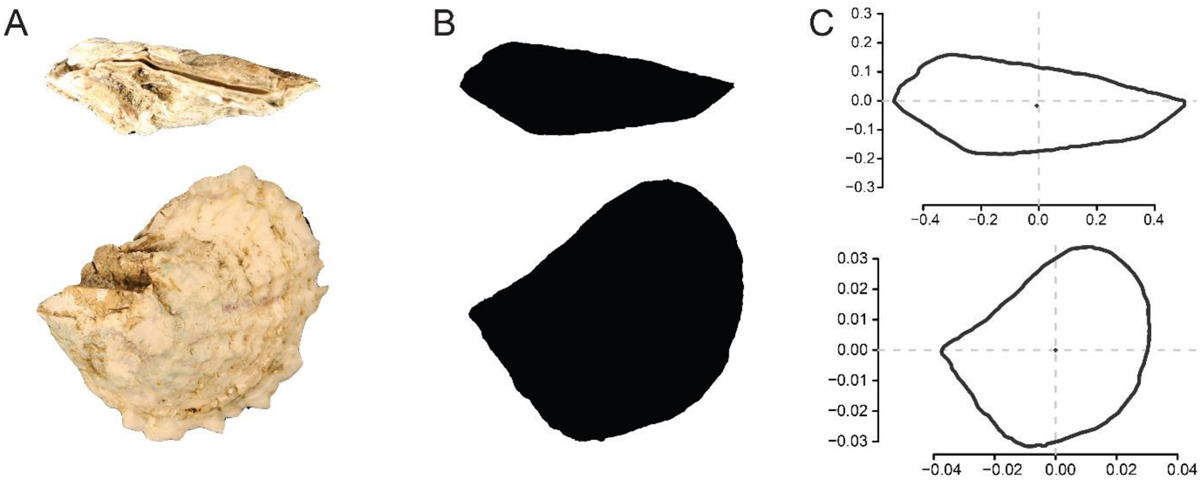

**Figure S1.** Digitization steps of oyster shell outlines: from digital photos to list of coordinates. **A.** High-resolution digital images of the shell, both lateral (top) and ventral (bottom) shell views, are acquired with a digital camera. **B.** Photographs are converted into black masks on a white background. **C.** Individual outlines are isolated, smoothed, and converted into a list of (*x*, *y*) pixel coordinates; coordinates are then used as input data for EFA.

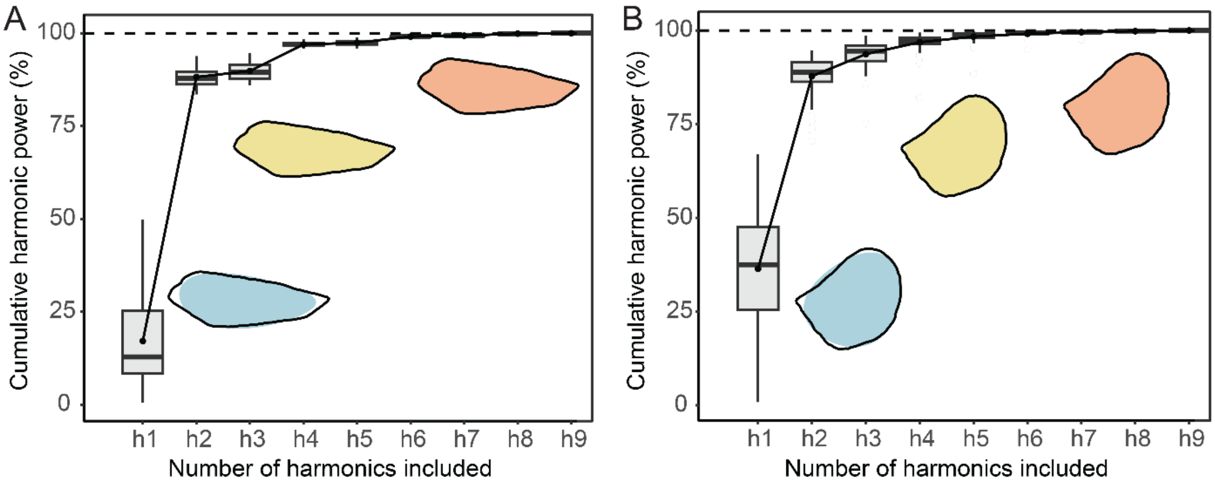

**Figure S2.** Calibration methods used in EFA and the inspection of principal components (PCs). Panels **A.** and **B.** show the cumulative spectrum of harmonic Fourier power for lateral and ventral shell views, respectively. Harmonic power is proportional to the harmonic amplitude and is considered as a measure of shape information. We retained seven harmonics because their cumulative power gathered 99% of the total cumulative power (Crampton, 1995). The average shell outline of *C. virginica* reconstructed with an increasing number of harmonics (3 - blue, 5 – yellow, and 8 - orange) is also displayed as a shaded area and compared with the original outlines (solid black line). A satisfactory reconstruction of shell outlines was achieved with six harmonics, whereas for nine harmonics, the approximation was almost perfect.

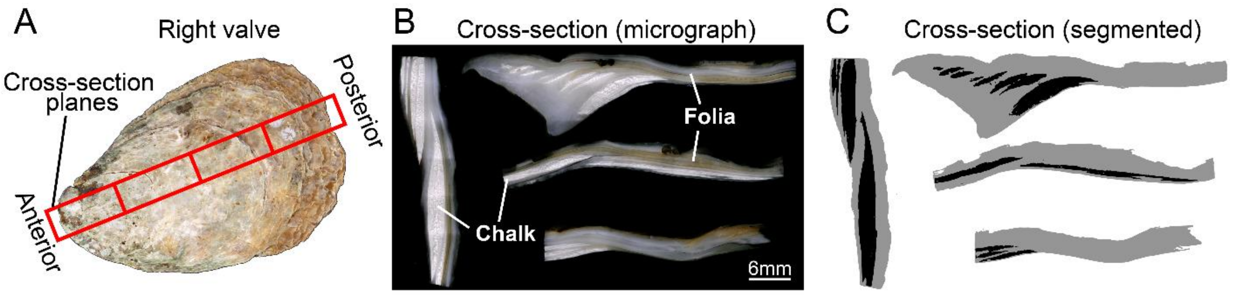

**Figure S3.** Protocol used for shell macro-structure analysis. (**A**) Shell tiles are sectioned along the axis of maximum growth. **B.** Micrographs of polished cross-sections are acquired with a digital microscope. **C.** Foliated (gray) and chalk (black) microstructure areas are segmented using the ilastik software (Berg *et al.*, 2019).

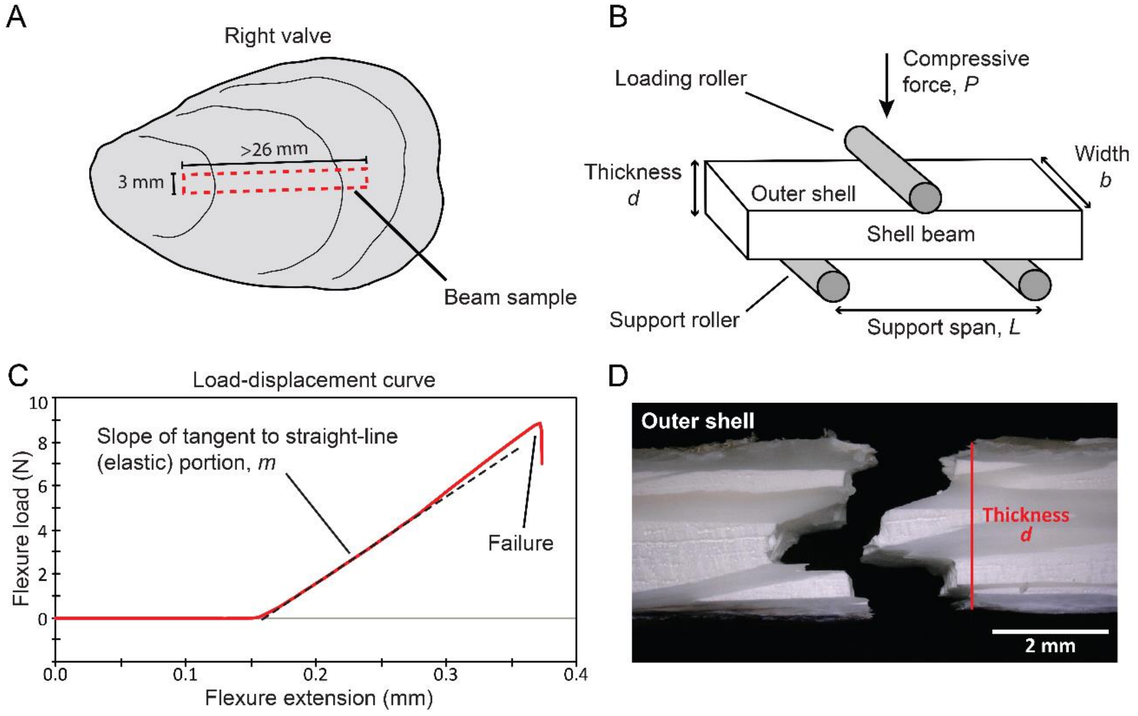

**Figure S4. A.** Beam position with respect to the shell of the specimen being tested is shown as a red dashed line. **B.** Diagram of three-point bending setup parameters. The span of the two supporting rollers was adjusted for each sample to obtain an acceptable span-to-thickness ratio (i.e., 16:1 or higher) to estimate precise material properties (ASTM, 2018). **C**. Typical load-displacement curve for a brittle material (oyster shell beam) which undergoes failure, along with tangent slope of the initial linear (elastic) loading portion. The Young's Modulus, *E*, is calculated using the sample width, *b*, the thickness at the fracture point, *d*, the distance between the two support rollers, *L*, and a force constant, *m*, that corresponds to the slope of the tangent to the initial (steepest) straight-line portion of the force-deflection curve for the tested sample:

$$E = 0.25\frac{L^3 m}{bd^3}$$

**D.** Example of fractographic inspection, showing the fracture pattern and thickness of the tested shell beam.

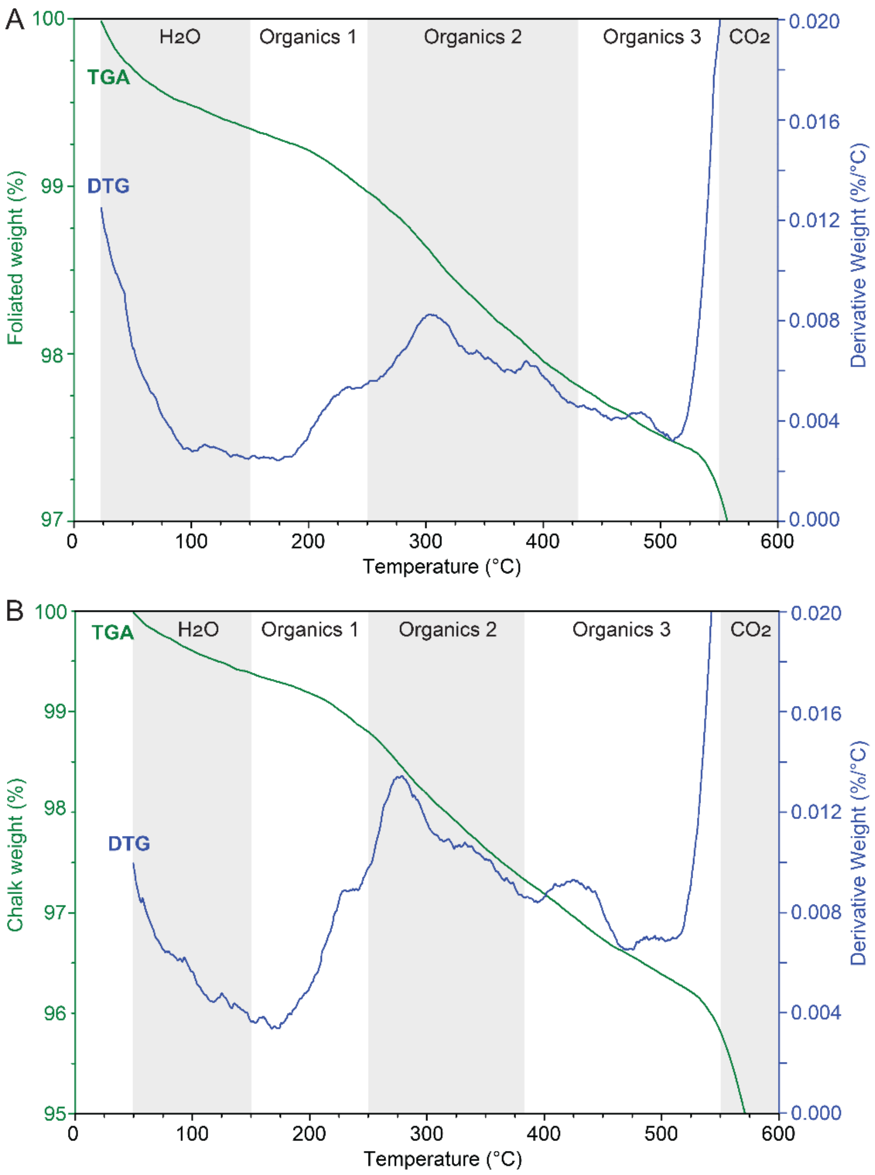

**Figure S5.** Example of thermal gravimetric analysis (TGA, green line) and derivative thermogravimetry (DTG, blue line) curves. The TGA curve shows how the weight changes with increasing temperature for the **A.** foliated and **B.** chalky calcite of *C. virginica*. The sample was heated at a linear rate of 10°C/min from ~25°C to 600°C. Five known regions of weight loss with increasing temperature are highlighted (Zaremba *et al.*, 1998; Harper, Checa & Rodríguez-Navarro, 2009): *i*) the evaporation of physically adsorbed water below 150°C, *ii*) the degradation of the non-proteinaceous component of the inter-crystalline organic matrix at 150-250°C, *iii*) the degradation of the proteinaceous component of the inter-crystalline organic matrix at 250-430°C for folia and 250-380°C for chalk, *iv*) the degradation of the intracrystalline organic matrix at 430-550°C for folia and 380-550°C for chalk, *v*) decomposition of calcium carbonate ($CaCO_3$) into calcium oxide (CaO) and carbon dioxide ($CO_2$) at temperature exceeding 550°C. The DTG curve corresponds to the derivative of the thermal curve and shows the rate of weight loss during heating. The peaks in the DTG curve represent the temperatures at which the organic mass loss is most rapid.

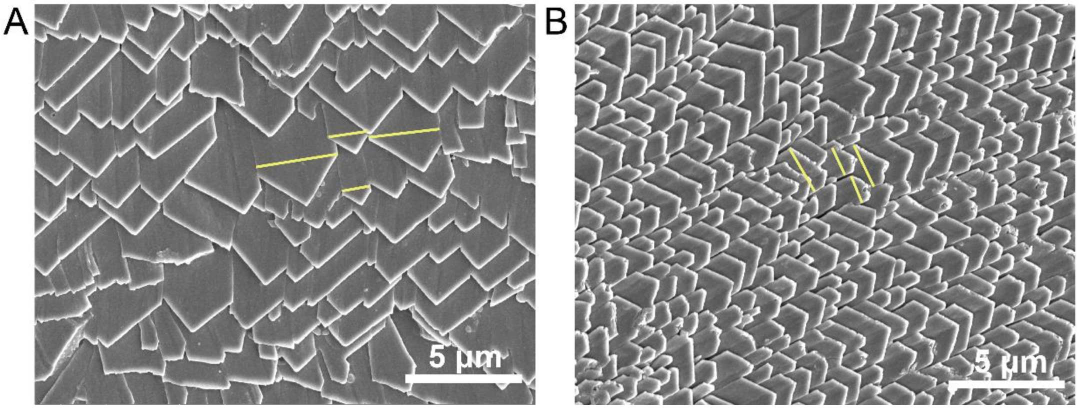

**Figure S6.** Example of SEM micrographs of oyster shell folia from **A.** LCL and **B.** MCB locations that were used to measure laths size (yellow lines), variability, and density. Pictures acquired with a FEI NovaNano 450.

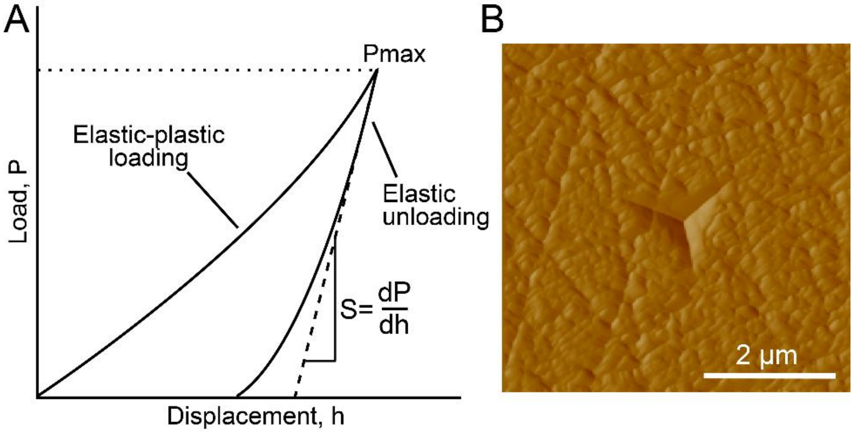

**Figure S7. A.** Example of loading-unloading curve adapted from Oliver & Pharr (2004) (Schabenberger & Pierce, 2002) showing indentation parameters. The reduced modulus, $E_r$, is calculated as

$$E_r = \frac{\sqrt{\pi}S}{2\beta\sqrt{A}}$$

where $\beta$ is a constant that depends on the geometry of the indenter, $S$ the slope of the initial portion of the unloading curve, and $A$ is the contact area of the indenter's tip. The effective modulus, $E$, is calculated from

$$\frac{1}{E_r} = \frac{1-\nu^2}{E} + \frac{1-\nu_i^2}{E_i}$$

where $E$ and $\nu$ are the specimen's effective modulus and Poisson's ratio respectively, and $E_i$ and $\nu_i$ are those of the indenter. The hardness, $H$, is determined as

$$H = \frac{P_{Max}}{A}$$

where $P_{max}$ is the maximum load. **B.** SPM map of single indent on foliated structure showing geometry of the Berkovich tip.

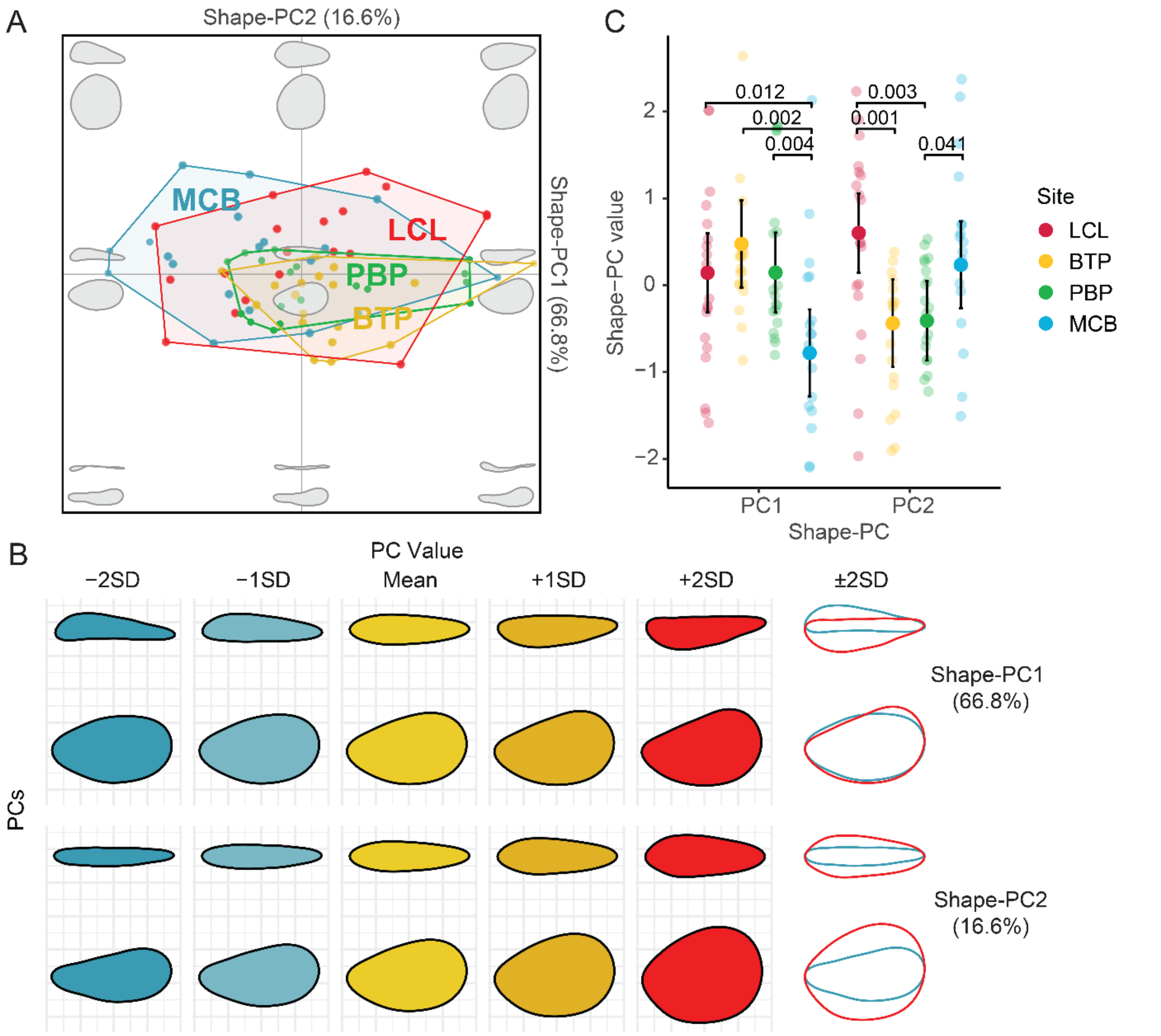

**Figure S8. A.** Scatterplot of the first two principal components (Shape-PCs) from a PCA performed on elliptic Fourier coefficients from an EFA, showing shape variability of orthogonal views of oyster shells from the HRE (location names as in Figure 1) among locations across the morphospace (reconstructed shapes on the background: lateral profile = top outline, ventral profile = bottom outline). Convex hulls (continuous lines) and the proportion of variance explained by each PC are reported. **B.** Between locations pairwise comparisons of Shape-PC values. P-values are reported only when differences between sites are significant. **C.** Contribution of the first two Shape-PCs to shape variation. The average shell shapes for lateral and ventral profiles were represented for increasing values along each Shape-PC and compared (red: Mean value +2SD; blue: Mean value –2SD). Shape-PC1 (66.8%) captured variations in the symmetry of the lateral profile (flat vs concave valves) and symmetry of the ventral profile along the shell growth axis (more vs less elliptic). Increasing Shape-PC1 values described flatter right valves, concave left valves, less elliptical/symmetric ventral profiles. Shape-PC2 (16.6%) captured variations in the width of the lateral profile and roundness of the ventral profile (elongated vs round). Increasing Shape-PC2 described wider lateral profiles and rounder ventral profiles.

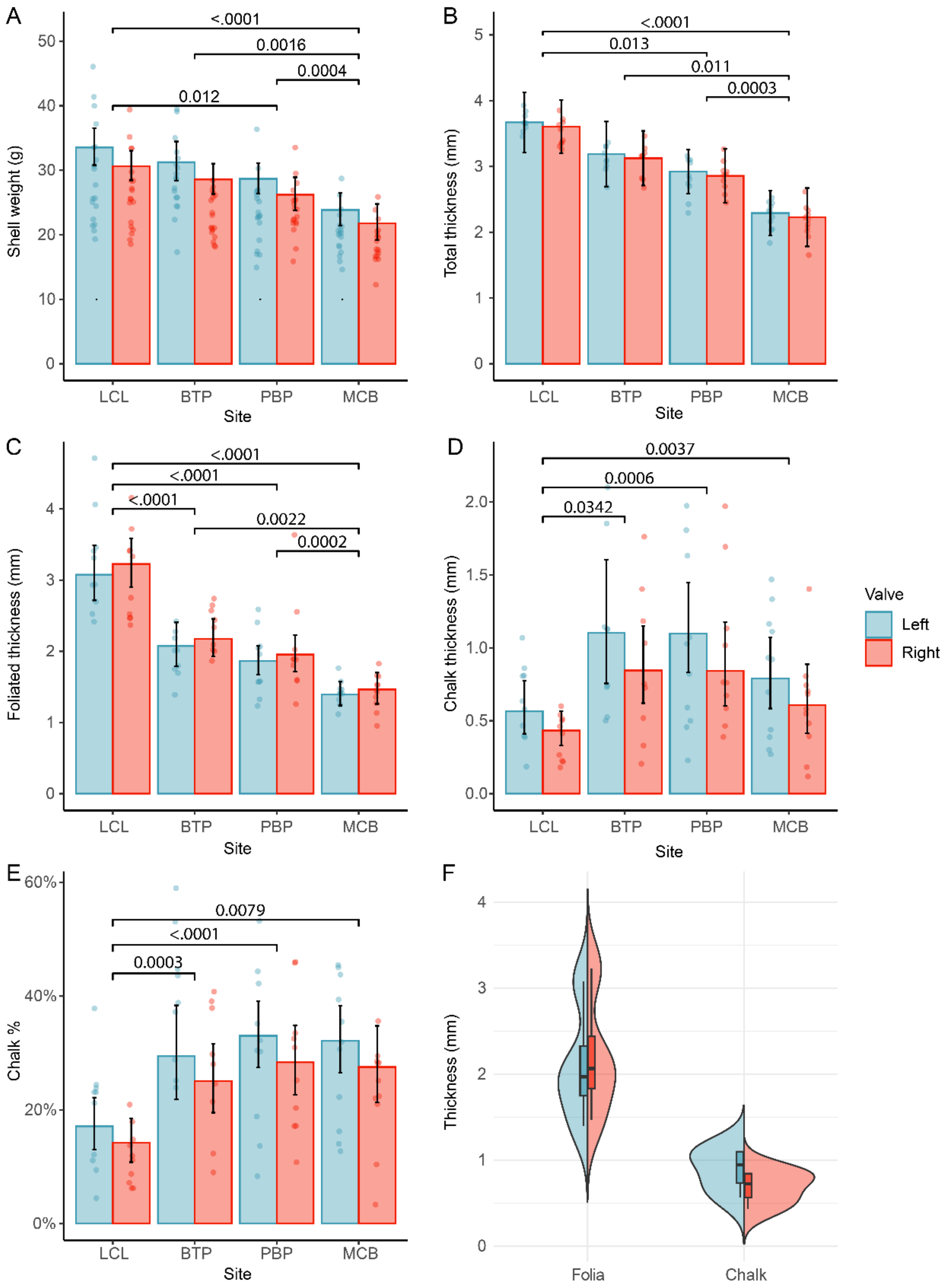

**Figure S9.** Variations in oyster **A.** shell weight, **B.** total shell thickness, **C.** folia thickness, **D.** chalk thickness, and **E.** proportion of chalk for both left (blue bars) and right (red bars) shell valves at different collection sites, as in Figure 1. The error bars represent 95% CIs. Significant pairwise contrasts between sites are reported as top brackets along with significant *P*-values. (**F**) Violin plot of pooled observations showing differences in folia and chalk thickness between left and right valves in the study region.

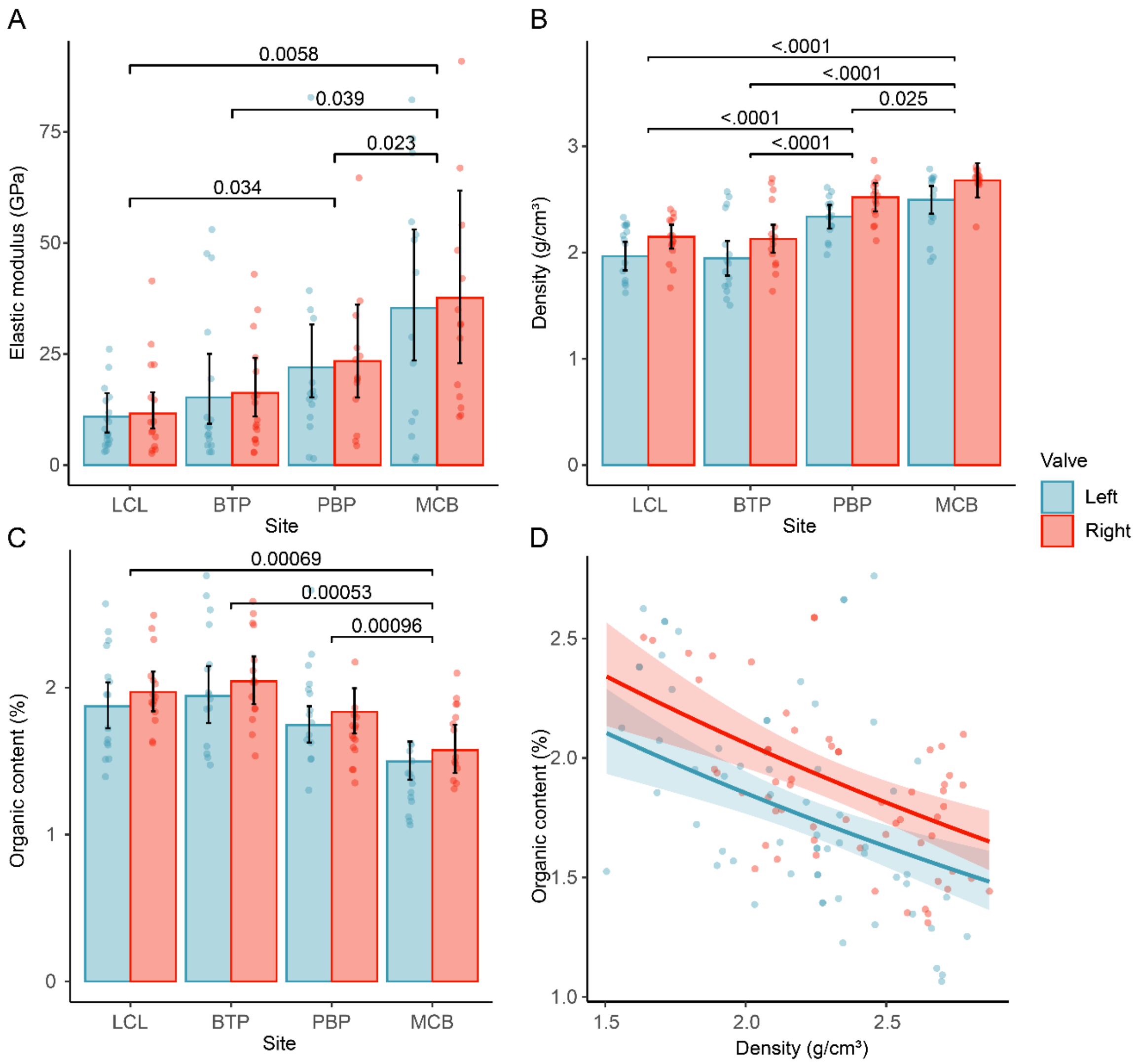

**Figure S10.** Variations in oyster shell **A.** elastic modulus, **B.** density, and **C.** organic content for both left (blue bars) and right (red bars) shell valves at different collection sites, as in Figure 1. The error bars represent 95% CIs. Significant pairwise contrasts between sites are reported as top brackets along with significant *P*-values. **D.** Relationship between shell organic content and density between left and right valves.

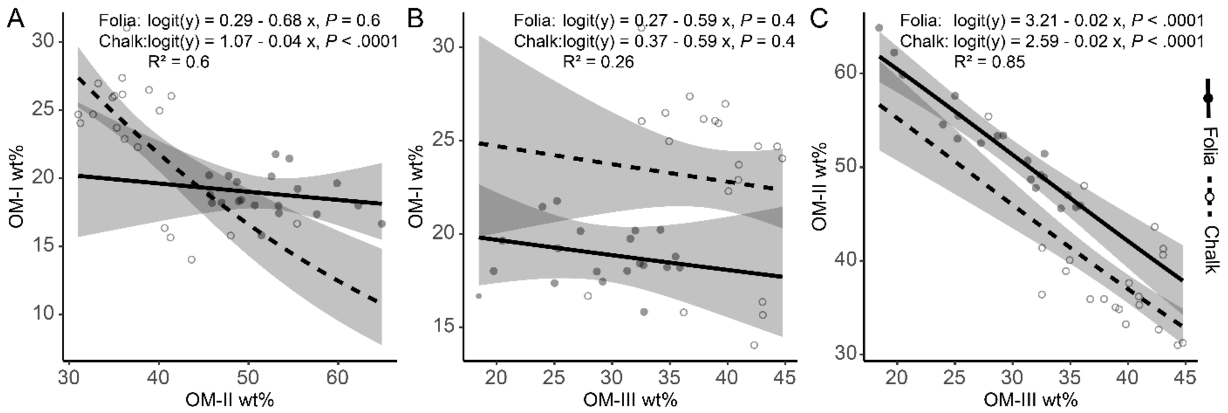

**Figure S11.** Predicted relationships between the relative wt% of OM component for foliated (solid line and filled dots) and chalky (dashed line and open circles) calcite: **A.** OM-I and OM-II, **B.** OM-I and OM-III, **C.** OM-II and OM-III. Shaded areas represent 95% CIs. Coefficients are reported on the logit-scale.

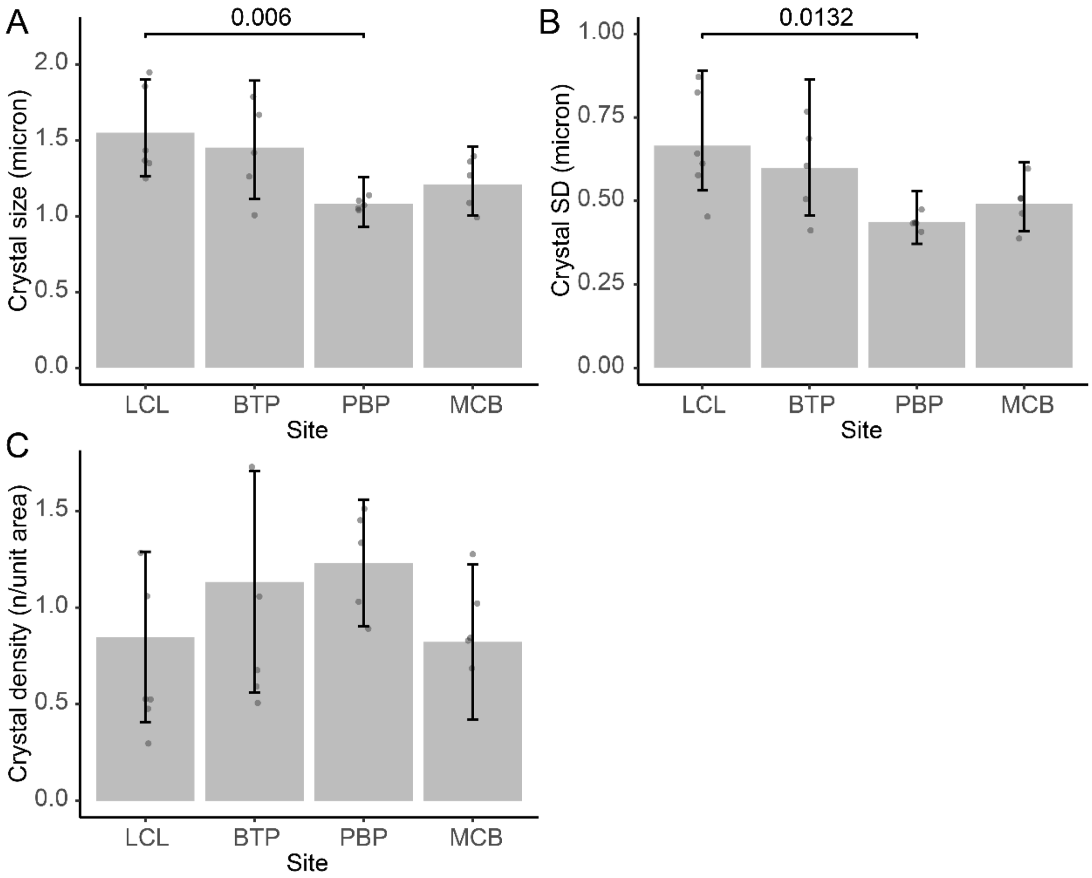

**Figure S12.** Variations in oyster shell calcitic laths (crystals) **A.** size, **B.** SD, and **C.** density at different collection sites. The error bars represent 95% CIs. Significant pairwise contrasts between sites are reported as top brackets along with significant *P*-values.

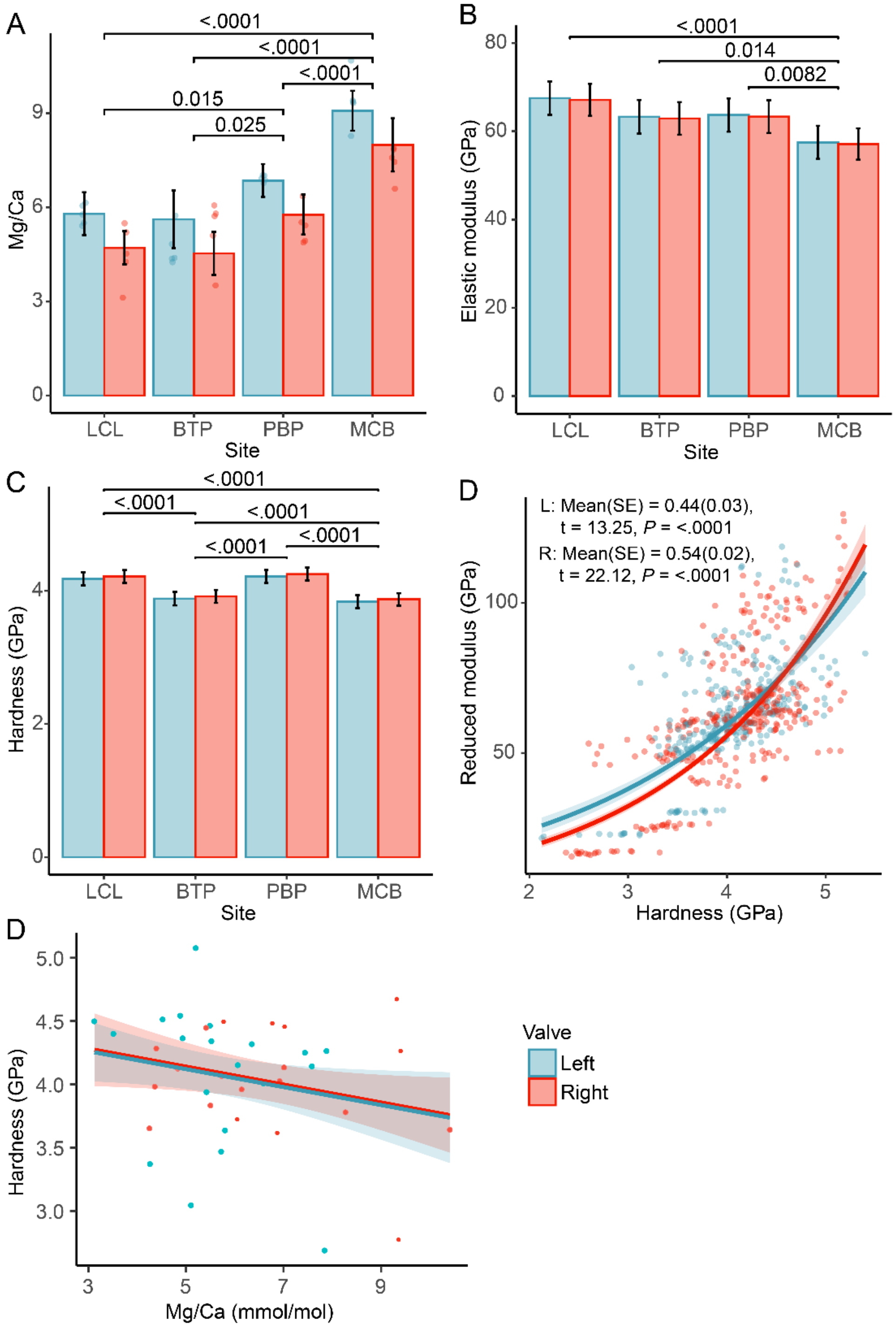

**Figure S13.** Variations in oyster shell **A.** Mg/Ca ratio, **B.** elastic modulus, and **C.** hardness for both left (blue bars) and right (red bars) shell valves at different collection sites. The error bars represent 95% CIs. Significant pairwise contrasts between sites are reported as top brackets along with significant *P*-values. **D.** Relationship between shell elastic modulus and hardness between left and right valves. **E.** Relationship between hardness and Mg/Ca ratio between left and right valves.

SUPPLEMENTARY TABLES

**Table S1.** Collection details for *Crassostrea virginica* shells used for this study. For each sampling location the site ID (as in Fig. 1), site coordinates (latitude and longitude), sample size (*n*), restorations method used, first install date, average sample shell height at install, and organization conducting restoration are reported. All the analyzed oyster specimens were collected from the subtidal zone at a depth of 0.5 m.

| **Location** | **ID** | **Latitude** | **Longitude** | ***n*** | **Restoration method** | **Install date** | **Size at install** | **Organization** |
|---|---|---|---|---|---|---|---|---|
| Mario M. Cuomo Bridge | MCB | 41°3'0.03"N | 73°53'49.49"W | 18 | Metal gabions | July 2018 | NA | BOP & HRF |
| Bush Terminal Park | BTP | 40°39'14.4"N | 74°01'08.5"W | 18 | Metal filing cabinet | August 2018 | ~5mm | BOP |
| Paerdegat Basin Park | PBP | 40°37'35.4"N | 73°54'15.9"W | 20 | Metal filing cabinet | July 2018 | ~5mm | BOP |
| Lemon Creek Lagoon | LCL | 40°30'44.0"N | 74°11'57.8"W | 19 | Metal gabions | August 2018 | ~5mm | BOP |

**Table S2.** Summary statistics for pairwise comparisons of Shape-PC1 and Shape-PC2 values sampling locations. Contrast values are the difference between locations' mean values. * denotes statistical significance.

| Shape-PC | Location | Contrast | SE | t-value | *P*-value | |
|---|---|---|---|---|---|---|
| Shape-PC1 | LCL-BTP | -0.333 | 0.314 | -1.059 | 0.29 | |
| | LCL-PBP | -0.003 | 0.340 | -0.008 | 0.99 | |
| | LCL-MCB | 0.924 | 0.362 | 2.549 | 0.012 | * |
| | BTP-PBP | 0.330 | 0.366 | 0.902 | 0.37 | |
| | BTP-MCB | 1.257 | 0.392 | 3.207 | 0.002 | * |
| | PB-MCB | 0.927 | 0.313 | 2.956 | 0.004 | * |
| Shape-PC2 | LCL-BTP | 1.041 | 0.314 | 3.31 | 0.001 | * |
| | LCL-PBP | 1.011 | 0.340 | 2.977 | 0.003 | * |
| | LCL-MCB | 0.367 | 0.362 | 1.012 | 0.31 | |
| | BTP-PBP | -0.029 | 0.366 | -0.08 | 0.94 | |
| | BTP-MCB | -0.674 | 0.392 | -1.72 | 0.09 | |
| | PB-MCB | -0.644 | 0.313 | -2.056 | 0.041 | * |

**Table S3.** GLMs summary statistics for *C. virginica* shell weight and deposition for both shell valves. Variation of shell weight ($n$ = 74 × valve), total thickness ($n$ = 40 × valve), foliated thickness ($n$ = 40 × valve), chalk thickness ($n$ = 40 × valve) and chalk proportion ($n$ = 40 × valve) were modeled as a function of collection location (Site), shell valve (Valve) and shell size (Area). Estimated statistics and significance values for the modelled relationships and pairwise comparisons among sampling locations are reported (Supplementary Material Fig. S9). * denotes statistical significance.

| **Model output** | | | | | **Pairwise comparisons** | | | | |
|---|---|---|---|---|---|---|---|---|---|
| Parameters | Estimates | SE | t-value | *P*-value | Locations | Contrast | SE | t-value | *P*-value |
| **Shell weight** | | | | | | | | | |
| (Intercept)$^{+}$ | -2.110 | 0.292 | -7.22 | <.0001 * | LCL/BTP | 1.070 | 0.051 | 1.48 | 0.14 |
| ln(Area) | 1.434 | 0.068 | 20.97 | <.0001 * | LCL/PB | 1.170 | 0.072 | 2.55 | 0.012 * |
| Site(BTP) | -0.070 | 0.048 | -1.48 | 0.14 | LCL/MCE | 1.410 | 0.107 | 4.49 | <.0001 * |
| Site(PBP) | -0.157 | 0.061 | -2.55 | 0.012 * | BTP/PB | 1.090 | 0.075 | 1.26 | 0.21 |
| Site(MCB) | -0.341 | 0.076 | -4.49 | <.0001 * | BTP/MCE | 1.310 | 0.110 | 3.23 | 0.0016 * |
| Valve(R) | -0.090 | 0.036 | -2.51 | 0.013 * | PB/MCB | 1.200 | 0.061 | 3.65 | 0.0004 * |
| **Total thickness** | | | | | | | | | |
| (Intercept)$^{+}$ | 2.835 | 0.529 | 5.36 | <.0001 * | LCL-BTP | 0.482 | 0.264 | 1.82 | 0.0723 |
| Area | 0.017 | 0.008 | 2.20 | 0.0313 * | LCL-PB | 0.749 | 0.295 | 2.54 | 0.0132 * |
| Site(BTP) | -0.482 | 0.264 | -1.82 | 0.0723 | LCL-MCE | 1.379 | 0.314 | 4.39 | <.0001 * |
| Site(PBP) | -0.749 | 0.295 | -2.54 | 0.0132 * | BTP-PB | 0.267 | 0.320 | 0.83 | 0.4067 |
| Site(MCB) | -1.379 | 0.314 | -4.39 | <.0001 * | BTP-MCE | 0.897 | 0.344 | 2.60 | 0.0111 * |
| Valve(R) | -0.063 | 0.147 | -0.43 | 0.6673 | PB-MCB | 0.630 | 0.167 | 3.78 | 0.0003 * |
| **Foliated thickness** | | | | | | | | | |
| (Intercept)$^{+}$ | 1.008 | 0.151 | 6.65 | <.0001 * | LCL/BTP | 1.480 | 0.102 | 5.72 | <.0001 * |
| Area | 0.002 | 0.002 | 1.09 | 0.28 | LCL/PB | 1.650 | 0.139 | 5.95 | <.0001 * |
| Site(BTP) | -0.394 | 0.069 | -5.72 | <.0001 * | LCL/MCE | 2.200 | 0.206 | 8.45 | <.0001 * |
| Site(PBP) | -0.501 | 0.084 | -5.95 | <.0001 * | BTP/PB | 1.110 | 0.107 | 1.11 | 0.68 |
| Site(MCB) | -0.790 | 0.093 | -8.45 | <.0001 * | BTP/MCE | 1.490 | 0.158 | 3.71 | 0.0022 * |
| Valve(R) | 0.047 | 0.052 | 0.92 | 0.36 | PB/MCB | 1.340 | 0.088 | 4.41 | 0.0002 * |
| **Chalk thickness** | | | | | | | | | |
| (Intercept)$^{+}$ | -0.085 | 0.254 | -0.34 | 0.74 | LCL-BTP | -0.549 | 0.155 | -3.55 | 0.0037 * |
| Area | 0.012 | 0.004 | 3.03 | 0.0034 * | LCL-PB | -0.540 | 0.132 | -4.09 | 0.0006 * |
| Site(BTP) | 0.549 | 0.155 | 3.55 | 0.0007 * | LCL-MCE | -0.358 | 0.129 | -2.78 | 0.034 * |
| Site(PBP) | 0.540 | 0.132 | 4.09 | 0.0001 * | BTP-PB | 0.009 | 0.203 | 0.04 | 0.98 |
| Site(MCB) | 0.358 | 0.129 | 2.78 | 0.0069 * | BTP-MCE | 0.191 | 0.204 | 0.94 | 0.79 |
| Valve(R) | -0.147 | 0.089 | -1.66 | 0.10 | PB-MCB | 0.183 | 0.108 | 1.69 | 0.34 |
| **Chalk %** | | | | | | | | | |
| (Intercept)$^{+}$ | -2.497 | 0.464 | -5.39 | <.0001 * | LCL-BTP | -0.116 | 0.036 | -3.18 | 0.0079 * |
| Area | 0.010 | 0.004 | 2.39 | 0.017 * | LCL-PB | -0.150 | 0.033 | -4.61 | <.0001 * |
| Site(BTP) | 0.703 | 0.208 | 3.37 | 0.0007 * | LCL-MCE | -0.142 | 0.035 | -4.03 | 0.0003 * |
| Site(PBP) | 0.870 | 0.191 | 4.56 | <.0001 * | BTP-PB | -0.034 | 0.046 | -0.75 | 0.88 |
| Site(MCB) | 0.829 | 0.204 | 4.07 | <.0001 * | BTP-MCE | -0.026 | 0.051 | -0.50 | 0.96 |
| Valve(R) | -0.219 | 0.139 | -1.58 | 0.11 | PB-MCB | 0.009 | 0.036 | 0.24 | 0.99 |

+Site LCL and Left valve are used as reference levels.

**Table S4.** GLMs summary statistics for *C. virginica* shell elasticity, density, and organic content for both shell valves. Variation of elasticity modulus ($n$ = 61 × valve), shell density ($n$ = 61 × valve), and organic content ($n$ = 61 × valve) were modeled as a function of collection location (Site), shell valve (Valve) and shell size (Area). Estimated statistics and significance values for the modelled relationships and pairwise comparisons among sampling locations are reported (see also Supplementary Material Figure S9). * denotes statistical significance.

| **Model output** | | | | | | **Pairwise comparisons** | | | | | |
|---|---|---|---|---|---|---|---|---|---|---|---|
| Parameters | Estimates | SE | t-value | *P*-value | | Locations | Contrast | SE | t-value | *P*-value | |
| **Modulus elasticity** | | | | | | | | | | | |
| (Intercept)$^{+}$ | 2.36 | 0.46 | 5.11 | <0.0001 | * | LCL/BTP | 1.02 | 0.15 | 0.12 | 0.90 | |
| Area | 0.00 | 0.01 | 0.09 | 0.93 | | LCL/PB | 0.82 | 0.15 | -1.06 | 0.29 | |
| Site(BTP) | 0.34 | 0.21 | 1.60 | 0.11 | | LCL/MCB | 0.63 | 0.13 | -2.22 | 0.028 | * |
| Site(PBP) | 0.70 | 0.28 | 2.53 | 0.013 | * | BTP/PB | 0.81 | 0.18 | -0.99 | 0.32 | |
| Site(MCB) | 1.18 | 0.31 | 3.76 | 0.0003 | * | BTP/MCB | 0.61 | 0.15 | -1.99 | 0.049 | * |
| Valve(R) | 0.06 | 0.16 | 0.39 | 0.70 | | PB/MCB | 0.76 | 0.12 | -1.76 | 0.08 | |
| **Shell density** | | | | | | | | | | | |
| (Intercept)$^{+}$ | 1.91 | 0.16 | 12.14 | < .0001 | * | LCL-BTP | 0.02 | 0.07 | 0.28 | 0.78 | |
| Area | 0.00 | 0.00 | 0.54 | 0.59 | | LCL-PB | -0.37 | 0.09 | -4.11 | 0.0001 | * |
| Site(BTP) | -0.02 | 0.07 | -0.28 | 0.78 | | LCL-MCB | -0.53 | 0.10 | -5.07 | <.0001 | * |
| Site(PBP) | 0.37 | 0.09 | 4.11 | <.0001 | * | BTP-PB | -0.39 | 0.10 | -3.73 | 0.0003 | * |
| Site(MCB) | 0.53 | 0.10 | 5.07 | <.0001 | * | BTP-MCB | -0.55 | 0.12 | -4.57 | <.0001 | * |
| Valve(R) | 0.18 | 0.05 | 3.50 | 0.0007 | * | PB-MCB | -0.16 | 0.07 | -2.28 | 0.025 | * |
| **Shell organic content** | | | | | | | | | | | |
| (Intercept)$^{+}$ | -4.007 | 0.099 | -40.68 | < .0001 | * | LCL-BTP | -0.0007 | 0.0008 | -0.88 | 0.38 | |
| Area | 0.001 | 0.001 | 0.73 | 0.46 | | LCL-PB | 0.0013 | 0.0010 | 1.26 | 0.21 | |
| Site(BTP) | 0.037 | 0.042 | 0.88 | 0.38 | | LCL-MCB | 0.0039 | 0.0011 | 3.44 | 0.0006 | * |
| Site(PBP) | -0.072 | 0.057 | -1.26 | 0.21 | | BTP-PB | 0.0020 | 0.0012 | 1.65 | 0.10 | |
| Site(MCB) | -0.228 | 0.067 | -3.39 | 0.0007 | * | BTP-MCB | 0.0046 | 0.0013 | 3.46 | 0.0005 | * |
| Valve(R) | 0.051 | 0.034 | 1.53 | 0.13 | | PB-MCB | 0.0025 | 0.0008 | 3.37 | 0.0008 | * |

+Site LCL and Left valve are used as reference levels.

**Table S5.** GLMs summary statistics for the modelled relationships: shell organic content and density ($n$ = 61 × valve), shell elastic modulus, density and organic content ($n$ = 61 × valve), with shell valve (Valve) and shell size (Area). * denotes statistical significance.

| Parameters | Estimates | SE | t-value | *P*-value | |
|---|---|---|---|---|---|
| **Organic content ~ Density** | | | | | |
| (Intercept)$^{+}$ | -3.580 | 0.147 | -24.33 | <.0001 | * |
| Area | 0.003 | 0.001 | 3.31 | 0.0009 | * |
| Density | -0.262 | 0.054 | -4.85 | <.0001 | * |
| Valve(Right) | 0.109 | 0.032 | 3.45 | 0.0006 | * |
| **Elastic modulus ~ Density + Organic content** | | | | | |
| (Intercept)$^{+}$ | -6.095 | 1.783 | -3.42 | 0.0009 | * |
| Area | 0.072 | 0.022 | 3.20 | 0.0018 | * |
| Density | 3.901 | 0.714 | 5.46 | <.0001 | * |
| Organics | -19.656 | 22.313 | -0.88 | 0.38 | |
| Valve(Right) | 2.444 | 1.068 | 2.29 | 0.024 | * |
| Density:Valve(Right) | -1.096 | 0.467 | -2.35 | 0.021 | * |
| Area:Density | -0.030 | 0.010 | -3.14 | 0.0021 | * |
| **Chalk% ~ Density** | | | | | |
| (Intercept)$^{+}$ | 3.086 | 0.851 | 3.63 | 0.0003 | * |
| Density | -2.145 | 0.451 | -4.76 | <.0001 | * |
| Area | 0.005 | 0.003 | 1.46 | 0.14 | |
| Valve(Right) | 0.343 | 0.211 | 1.63 | 0.10 | |

+Left valve is used as reference levels.

**Table S6**. GLMs summary statistics for the modeled relationships between the OM bulk wt% (*n* = 40) and the OM components wt% (*n* = 60 × valve) between sampling locations (Site) and between Layers (Folia, Chalk).* denotes statistical significance.

**Model output**

| Parameters | Estimates | SE | z-value | *P* -value | |
|---|---|---|---|---|---|
| **Bulk OM wt%** | | | | | |
| (Intercept)[+] | -3.86 | 0.07 | -56.63 | <.0001 | * |
| Site(BTP) | -0.15 | 0.10 | -1.46 | 0.15 | |
| Site(PBP) | -0.01 | 0.10 | -0.12 | 0.90 | |
| Site(MCB) | -0.27 | 0.10 | -2.62 | 0.0089 | * |
| Layer(Chalk) | 0.41 | 0.09 | 4.66 | <.0001 | * |
| Site(BTP):Layer(Chalk) | 0.18 | 0.13 | 1.38 | 0.17 | |
| Site(PBP):Layer(Chalk) | 0.02 | 0.12 | 0.13 | 0.89 | |
| Site(MCB):Layer(Chalk) | 0.35 | 0.13 | 2.70 | 0.0069 | * |
| **Components OM wt%** | | | | | |
| (Intercept)[‡] | -5.56 | 0.11 | -49.15 | <.0001 | * |
| Site(BTP) | -0.04 | 0.16 | -0.22 | 0.83 | |
| Site(PBP) | -0.03 | 0.16 | -0.16 | 0.87 | |
| Site(MCB) | -0.23 | 0.17 | -1.34 | 0.18 | |
| Layer(Chalk) | 0.73 | 0.14 | 5.31 | <.0001 | * |
| TreatmentTreatment2 | 1.08 | 0.13 | 8.19 | <.0001 | * |
| TreatmentTreatment3 | 0.40 | 0.15 | 2.74 | 0.006 | * |
| Site(BTP):Layer(Chalk) | 0.04 | 0.20 | 0.19 | 0.85 | |
| Site(PBP):Layer(Chalk) | 0.05 | 0.20 | 0.26 | 0.80 | |
| Site(MCB):Layer(Chalk) | -0.18 | 0.21 | -0.85 | 0.40 | |
| Layer(Chalk):OM(OM-II) | -0.76 | 0.17 | -4.55 | <.0001 | * |
| Layer(Chalk):OM(OM-III) | 0.02 | 0.18 | 0.12 | 0.90 | |
| Site(BTP):OM(OM-II) | -0.19 | 0.19 | -0.98 | 0.33 | |
| Site(PBP):OM(OM-II) | 0.03 | 0.19 | 0.15 | 0.88 | |
| Site(MCB):OM(OM-II) | -0.12 | 0.20 | -0.60 | 0.55 | |
| Site(BTP):OM(OM-III) | -0.04 | 0.21 | -0.17 | 0.87 | |
| Site(PBP):OM(OM-III) | -0.03 | 0.21 | -0.16 | 0.88 | |
| Site(MCB):OM(OM-III) | 0.07 | 0.22 | 0.33 | 0.74 | |
| Site(BTP):Layer(Chalk):OM(OM-II) | 0.27 | 0.24 | 1.14 | 0.25 | |
| Site(PBP):Layer(Chalk):OM(OM-II) | -0.06 | 0.24 | -0.23 | 0.82 | |
| Site(MCB):Layer(Chalk):OM(OM-II | 0.87 | 0.25 | 3.45 | 0.001 | * |
| Site(BTP):Layer(Chalk):OM(OM-III | 0.03 | 0.25 | 0.11 | 0.91 | |
| Site(PBP):Layer(Chalk):OM(OM-III | 0.01 | 0.25 | 0.03 | 0.98 | |
| Site(MCB):Layer(Chalk):OM(OM-II | 0.38 | 0.27 | 1.44 | 0.15 | |

**Pairwise comparisons**

| Group | Contrast | Estimates | SE | z-value | *P* -value | |
|---|---|---|---|---|---|---|
| Folia | LCL-BTP(Folia) | 0.0028 | 0.0019 | 1.46 | 0.14 | |
| | LCL-PB(Folia) | 0.0002 | 0.0020 | 0.12 | 0.90 | |
| | LCL-MCB(Folia | 0.0048 | 0.0018 | 2.62 | 0.009 | * |
| | BTP-PB(Folia) | -0.0025 | 0.0019 | -1.34 | 0.18 | |
| | BTP-MCB(Folia | 0.0021 | 0.0018 | 1.17 | 0.24 | |
| | PB-MCB(Folia) | 0.0046 | 0.0018 | 2.50 | 0.012 | * |
| Chalk | LCL-BTP(Chalk | -0.0009 | 0.0024 | -0.38 | 0.70 | |
| | LCL-PB(Chalk) | -0.0001 | 0.0024 | -0.06 | 0.95 | |
| | LCL-MCB(Chall | -0.0025 | 0.0024 | -1.02 | 0.31 | |
| | BTP-PB(Chalk) | 0.0008 | 0.0024 | 0.32 | 0.75 | |
| | BTP-MCB(Chal | -0.0016 | 0.0024 | -0.64 | 0.52 | |
| | PB-MCB(Chalk) | -0.0023 | 0.0024 | -0.96 | 0.34 | |
| OM-I Folia | LCL - BTP | 0.0001 | 0.0006 | 0.22 | 0.83 | |
| | LCL - PB | 0.0001 | 0.0006 | 0.16 | 0.87 | |
| | LCL - MCB | 0.0008 | 0.0006 | 1.34 | 0.18 | |
| | BTP - PB | 0.0000 | 0.0006 | -0.06 | 0.95 | |
| | BTP - MCB | 0.0006 | 0.0006 | 1.13 | 0.26 | |
| | PB - MCB | 0.0007 | 0.0006 | 1.19 | 0.24 | |
| OM-II Folia | LCL - BTP | 0.0022 | 0.0010 | 2.21 | 0.027 | * |
| | LCL - PB | 0.0000 | 0.0011 | -0.02 | 0.99 | |
| | LCL - MCB | 0.0033 | 0.0010 | 3.34 | <.0001 | * |
| | BTP - PB | -0.0022 | 0.0010 | -2.23 | 0.026 | * |
| | BTP - MCB | 0.0010 | 0.0009 | 1.14 | 0.25 | |
| | PB - MCB | 0.0033 | 0.0010 | 3.36 | <.0001 | * |
| OM-III Folia | LCL - BTP | 0.0004 | 0.0007 | 0.52 | 0.60 | |
| | LCL - PB | 0.0003 | 0.0007 | 0.44 | 0.66 | |
| | LCL - MCB | 0.0008 | 0.0007 | 1.14 | 0.26 | |
| | BTP - PB | -0.0001 | 0.0007 | -0.09 | 0.93 | |
| | BTP - MCB | 0.0004 | 0.0007 | 0.61 | 0.54 | |
| | PB - MCB | 0.0005 | 0.0007 | 0.70 | 0.48 | |
| OM-I Chalk | LCL - BTP | 0.0000 | 0.0009 | -0.02 | 0.98 | |
| | LCL - PB | -0.0002 | 0.0009 | -0.22 | 0.82 | |
| | LCL - MCB | 0.0026 | 0.0008 | 3.26 | 0.001 | * |
| | BTP - PB | -0.0002 | 0.0009 | -0.20 | 0.84 | |
| | BTP - MCB | 0.0027 | 0.0008 | 3.28 | 0.001 | * |
| | PB - MCB | 0.0028 | 0.0008 | 3.48 | <.0001 | * |
| OM-II Chalk | LCL - BTP | -0.0010 | 0.0011 | -0.95 | 0.34 | |
| | LCL - PB | 0.0000 | 0.0010 | 0.04 | 0.97 | |
| | LCL - MCB | -0.0044 | 0.0011 | -3.83 | <.0001 | * |
| | BTP - PB | 0.0011 | 0.0011 | 0.99 | 0.32 | |
| | BTP - MCB | -0.0033 | 0.0012 | -2.89 | 0.004 | * |
| | PB - MCB | -0.0044 | 0.0011 | -3.87 | <.0001 | * |
| OM-III Chalk | LCL - BTP | 0.0001 | 0.0011 | 0.06 | 0.95 | |
| | LCL - PB | 0.0000 | 0.0011 | 0.01 | 1.00 | |
| | LCL - MCB | -0.0006 | 0.0011 | -0.56 | 0.58 | |
| | BTP - PB | -0.0001 | 0.0011 | -0.06 | 0.95 | |
| | BTP - MCB | -0.0007 | 0.0011 | -0.62 | 0.54 | |
| | PB - MCB | -0.0006 | 0.0011 | -0.56 | 0.58 | |

+Site(LCL) and Layer(Folia) are used as reference levels

‡Site(LCL), Layer(Folia) and OM(OM-I) are used as reference levels

**Table S7**. GLMs summary statistics for the modelled relationships between crystal size (*n* = 21), SD (*n* = 21), density (*n* = 21), sampling location (Site) and shell size (Area). * denotes statistical significance.

| **Model output** | | | | | | **Pairwise comparisons** | | | | | |
|---|---|---|---|---|---|---|---|---|---|---|---|
| Parameters | Estimates | SE | t-value | *P*-value | | Contrast | Estimates | SE | t-value | *P*-value | |
| **Crystal size** | | | | | | LCL/BTP | 1.067 | 0.115 | 2.10 | 0.55 | |
| (Intercept)$^+$ | 0.466 | 0.268 | 1.74 | 0.10 | | LCL/PB | 1.435 | 0.188 | 3.27 | 0.014 | * |
| Area | -0.001 | 0.004 | -0.15 | 0.88 | | LCL/MCB | 1.281 | 0.211 | 2.57 | 0.15 | |
| Site(BTP) | -0.065 | 0.108 | -0.61 | 0.55 | | BTP/PB | 1.345 | 0.211 | 0.85 | 0.08 | |
| Site(PBP) | -0.361 | 0.131 | -2.76 | 0.01 | * | BTP/MCB | 1.200 | 0.233 | 0.99 | 0.36 | |
| Site(MCB) | -0.247 | 0.165 | -1.50 | 0.15 | | PB/MCB | 0.892 | 0.104 | 0.54 | 0.34 | |
| **Crystal SD** | | | | | | LCL-BTP | -0.170 | 0.201 | -0.85 | 0.41 | |
| (Intercept)$^+$ | 1.489 | 0.519 | 2.87 | 0.011 | * | LCL-PB | -0.788 | 0.283 | -2.79 | 0.01 | * |
| Area | 0.000 | 0.007 | 0.04 | 0.97 | | LCL-MCB | -0.532 | 0.330 | -1.61 | 0.13 | |
| Site(BTP) | 0.170 | 0.201 | 0.85 | 0.41 | | BTP-PB | -0.618 | 0.334 | -1.85 | 0.08 | |
| Site(PBP) | 0.788 | 0.283 | 2.79 | 0.013 | * | BTP-MCB | -0.362 | 0.391 | -0.93 | 0.37 | |
| Site(MCB) | 0.532 | 0.330 | 1.61 | 0.13 | | PB-MCB | 0.256 | 0.283 | 0.91 | 0.38 | |
| **Crystal density** | | | | | | LCL-BTP | -0.286 | 0.233 | -1.40 | 0.2 | |
| (Intercept)$^+$ | 1.221 | 0.580 | 2.11 | 0.05 | | LCL-PB | -0.384 | 0.283 | -1.32 | 0.2 | |
| Area | -0.007 | 0.008 | -0.94 | 0.36 | | LCL-MCB | 0.025 | 0.356 | 0.12 | 0.9 | |
| Site(BTP) | 0.286 | 0.233 | 1.23 | 0.24 | | BTP-PB | -0.098 | 0.339 | -0.14 | 0.8 | |
| Site(PBP) | 0.384 | 0.283 | 1.36 | 0.19 | | BTP-MCB | 0.311 | 0.420 | 0.88 | 0.5 | |
| Site(MCB) | -0.025 | 0.356 | -0.07 | 0.94 | | PB-MCB | 0.409 | 0.253 | 1.65 | 0.1 | |

**Table S8**. GLMs summary statistics for the modelled relationships between Mg/Ca ratios (*n* = 40), elastic modulus (*n* = 40) and hardness (*n* = 40) with sampling location (Site), shell size (Area) and valve (Valve). * denotes statistical significance.

| Model output | | | | | | Pairwise Contrasts | | | | | |
|---|---|---|---|---|---|---|---|---|---|---|---|
| Parameters | Estimates | SE | t-value | *P*-value | | Contrast | Estimates | SE | t-value | *P*-value | |
| **Mg/Ca** | | | | | | | | | | | |
| (Intercept)$^{+}$ | 6.751 | 0.950 | 7.11 | <.0001 | * | LCL-BTP | 0.178 | 0.360 | 0.50 | 0.62 | |
| Area | -0.019 | 0.014 | -1.38 | 0.18 | | LCL-PB | -1.057 | 0.412 | -2.56 | 0.015 | |
| Site(BTP) | -0.178 | 0.360 | -0.50 | 0.62 | | LCL-MCB | -3.279 | 0.518 | -6.33 | <.0001 | * |
| Site(PBP) | 1.057 | 0.412 | 2.56 | 0.015 | * | BTP-PB | -1.236 | 0.527 | -2.35 | 0.025 | |
| Site(MCB) | 3.279 | 0.518 | 6.33 | <.0001 | * | BTP-MCB | -3.457 | 0.650 | -5.32 | <.0001 | |
| Valve(Right) | -3.191 | 0.691 | -4.62 | <.0001 | * | PB-MCB | -2.222 | 0.355 | -6.25 | <.0001 | * |
| Area:Valve(Right) | 0.042 | 0.013 | 3.28 | 0.0024 | * | | | | | | |
| **Elastic modulus** | | | | | | | | | | | |
| (Intercept)$^{+}$ | 67.51 | 1.90 | 35.52 | <.0001 | * | LCL-BTP | 4.227 | 2.370 | 1.79 | 0.075 | |
| Site(BTP) | -4.23 | 2.37 | -1.79 | 0.075 | | LCL-PB | 3.818 | 2.360 | 1.62 | 0.11 | |
| Site(PBP) | -3.82 | 2.36 | -1.62 | 0.11 | | LCL-MCB | 10.019 | 2.330 | 4.30 | <.0001 | * |
| Site(MCB) | -10.02 | 2.33 | -4.30 | <.0001 | * | BTP-PB | -0.409 | 2.370 | -0.17 | 0.86 | |
| Valve(Right) | -0.38 | 1.67 | -0.23 | 0.82 | | BTP-MCB | 5.792 | 2.340 | 2.48 | 0.014 | * |
| | | | | | | PB-MCB | 6.201 | 2.340 | 2.65 | 0.008 | * |
| **Harndess** | | | | | | | | | | | |
| (Intercept)$^{+}$ | 4.178 | 0.050 | 83.77 | <.0001 | * | LCL-BTP | 0.298 | 0.062 | 4.80 | <.0001 | * |
| Site(BTP) | -0.298 | 0.062 | -4.80 | <.0001 | * | LCL-PB | -0.036 | 0.062 | -0.58 | 0.57 | |
| Site(PBP) | 0.036 | 0.062 | 0.58 | 0.57 | | LCL-MCB | 0.343 | 0.061 | 5.61 | <.0001 | * |
| Site(MCB) | -0.343 | 0.061 | -5.61 | <.0001 | * | BTP-PB | -0.334 | 0.062 | -5.35 | <.0001 | * |
| Valve(Right) | 0.034 | 0.044 | 0.78 | 0.44 | | BTP-MCB | 0.045 | 0.061 | 0.73 | 0.46 | |
| | | | | | | PB-MCB | 0.379 | 0.061 | 6.17 | <.0001 | * |

+Site(LCL) andValve(Right) are used as reference levels

**Table S9.** Variable loadings on the first two principal components (PCs) from a PCA performed on environmental descriptors (Fig. 6A), macro scale (Fig. 6B) and micro scale (Fig. 6C) shell traits. The proportion of variance explained by each PC is reported. Bold values indicate the most influential variables in each PCA.

| **Enviro-PCA** | | | **Macro-PCA** | | | **Micro-PCA** | | |
|---|---|---|---|---|---|---|---|---|
| Parameter | PC1 74.85% | PC2 23.44% | Parameter | PC1 29.38% | PC2 25.77% | Parameter | PC1 31.06% | PC2 21.94% |
| Temperature | -0.030 | **0.561** | Total | **0.410** | -0.283 | Elastic modulus | 0.110 | **-0.264** |
| Salinity | -0.292 | **-0.463** | Foliated | **0.486** | 0.146 | Hardness | **0.205** | -0.026 |
| Chl-a | **-0.445** | 0.268 | Chalk | -0.007 | **-0.614** | Crystal size | 0.281 | **-0.390** |
| Turbidity | **-0.462** | 0.084 | Chalk % | -0.146 | **-0.617** | Crystal SD | 0.283 | **-0.378** |
| DO | **-0.470** | 0.065 | Shape-PC1 | **0.251** | 0.000 | Crystal density | -0.074 | **0.439** |
| pH | **-0.461** | -0.069 | Shape-PC2 | **0.369** | 0.228 | Mg/Ca | **-0.415** | -0.005 |
| | | | Elastic modulus | **-0.460** | 0.033 | OM-I wt%-FOL | **0.414** | 0.262 |
| | | | Organics% | **0.269** | -0.226 | OM-II wt%-FOL | **0.379** | 0.115 |
| | | | Density | **-0.303** | 0.191 | OM-III wt%-FOL | 0.173 | **0.436** |
| | | | | | | OM-I wt%-CHA | **0.360** | 0.174 |
| | | | | | | OM-II wt%-CHA | **-0.363** | 0.072 |
| | | | | | | OM-III wt%-CHA | 0.045 | **0.367** |

SUPPLEMENTARY DOCUMENT S1

Details from the Billion Oyster Project - NYC Oyster Monitoring Report 2018 (Burmester & McCann, 2018; Baumann, Burmester & Castro, 2021) and HRECOS (https://www.hrecos.org/) datasets used in this study.

**BOP Continuous Water Quality Measurements:**
Since 2017, during each field season (from May to November), the Billion Oyster Project has deployed Onset HOBO water quality loggers (U-24 and U-26) to record high-resolution (every 15-minutes) time series data of dissolved oxygen, salinity, and temperature. The loggers were monitored several times each year during deployment to remove fouling organisms and sediment build-up, offload data, and recalibrate by taking independent water quality measurements made with a Horiba U-52 multiparameter instrument. Onset HOBOware software was used to adjust the dissolved oxygen values based on the salinity time series and to account for fouling. Due to differences in deployment and maintenance schedules, and adherence to quality assurance protocols, some time periods were excluded, resulting in data that do not span precisely the same time periods across sites.

**BOP Point Water Quality Measurements:**
On several sampling dates, point water quality data were measured using a Horiba U-52 multiparameter water quality probe to measure temperature, dissolved oxygen, salinity, pH, conductivity, turbidity, and total dissolved solids. All samples were taken just below the water surface, adjacent to the restored oysters. Typically, three replicate samples were taken at least two minutes apart on each sampling date. These samples only provide a snapshot of the water quality conditions at a site and were taken in conjunction with the regular maintenance of Onset HOBO water quality data loggers. The Horiba U-52 can also measure parameters not recorded by the water quality data loggers, such as pH and turbidity.

**BOP Chlorophyll-*a* Measurements:**
At LCL, BTP, and PBP sites, chlorophyll-a concentration was measured from discrete water samples. Water samples were collected next to the reefs at regular intervals during the 2017-2018 monitoring field seasons. A known volume of water was filtered on a 0.7-micron glass fiber filter to measure available chlorophyll-*a*. Additional samples from the 5-20 µm size fraction were measured for chlorophyll-*a*, which is a proxy for food supply to our oysters (Newell & Langdon, 1996). Chlorophyll was extracted from the filters with 90% acetone and quantified via fluorometry with a Turner Designs AquaFluor handheld fluorometer in the BOP Hatchery on Governors Island. Chlorophyll-*a* values were corrected via acidification to exclude phaeopigments. On each sampling date, three water samples for each filter size were analyzed to quantify variation. Each site was sampled on three to five dates between June and October 2017-2018. These chlorophyll-*a* results are further supported by data collected on phytoplankton species identity and biochemical composition, and quantity.

**HRECOS Continuous Water Quality Measurements Station:**
The HRECOS continuous water quality monitoring data were obtained from the station located at the end of Piermont Pier, Piermont, NY (Station name: "Hudson River at Piermont NY"; HRECOS ID: HRPMNT; Site ID: 01376269). The system uses an EXO 6600 (YSI) multiparameter water

quality probe to collect continuous measurements of water temperature, salinity, conductivity, pH, dissolved oxygen, turbidity, and chlorophyll-*a* every 15 minutes since 2010. The station is maintained in partnership with the Lamont-Doherty Earth Observatory. Data can be accessed through: https://waterdata.usgs.gov/monitoring-location/01376269/#parameterCode=00010&period=P7D (last access date 05/02/2023).

SUPPLEMENTARY METHODS S1

### Geometric morphometrics analysis

The shape of *Crassostrea virginica* shells was analysed through an elliptic Fourier analysis (EFA) of outlines (Giardina & Kuhl, 1977; Kuhl & Giardina, 1982; Bonhomme *et al.*, 2014; Telesca *et al.*, 2018). This geometric morphometrics approach (Rohlf & Marcus, 1993; Adams, Rohlf & Slice, 2004) was performed in order to examine shell shape variation within and between groups of individuals. Unlike traditional morphometrics approaches, which consider outlines as a collection of composite ratios, EFA considers outlines as a whole, taking into account all the geometrical relationships of the input data.

Oyster morphology has been studied mostly using a range of composite ratios (Palmer & Carriker, 1979; Thomas, Allen & Plough, 2019; Hajovsky, Beseres Pollack & Anderson, 2021) that were originally developed as metrics to describe suitable shell shapes for the oyster market (Brake, Evans & Langdon, 2003). However, these derived variables are poor shape descriptors because they assume linear relationships between traits (Nakagawa *et al.*, 2017), can lead to spurious correlations (Kronmal, 1993), and different shapes can have the same shape ratios (Bonhomme, Frelat & Gaucherel, 2013). In contrast, the EFA uses Fourier transformations to extract geometric information from complex outlines (A. J. Haines & Crampton, 2000) and provides a more suitable approach to describe highly variable and complex oyster shapes.

EFA is a powerful method for extracting geometric information and has been implemented using the concept of Fourier series, which decomposes a periodic function into a sum of simpler trigonometric functions (such as sine and cosine) (J. Claude, 2008; Bonhomme *et al.*, 2014). These simple functions have frequencies that are integer multiples, which makes them harmonics of one another. This approach fits Fourier series separately on the x and y coordinates of an outline, projected on the Cartesian plane, as a function of the curvilinear abscissa (Kuhl & Giardina, 1982; J. Claude, 2008; Bonhomme *et al.*, 2014). EFA is then used to extract the geometrical information from outlines, described as periodic functions (Kuhl & Giardina, 1982; Rohlf & Archie, 1984), through their decomposition into the harmonic sum of trigonometric functions, called harmonics. Low-frequency harmonics approximate coarse-scale trends in the original outlines, while high-frequency harmonics fit their fine-scale variations (Bonhomme *et al.*, 2014). These can be normalized to remove homothetic, translational or rotational differences between shapes, and smoothed in order to remove outline noise during the digitization process or small outline irregularities (A. J. Haines & Crampton, 2000). Harmonic coefficients are then extracted and used as shape variables capturing shape information. The geometric information contained in the outlines is thus quantified and can be analysed with classical multivariate tools (such as multivariate analysis of variance, principal component analysis, and linear discriminant analysis).

EFA of outlines allows shape reconstruction from the numerical signature and this improved method has great advantages compared to more traditional approaches (A. J. Haines & Crampton, 2000; Adams, Rohlf & Slice, 2004; Bonhomme *et al.*, 2014; Telesca *et al.*, 2018): complex shapes can be fitted, outlines smoothed, starting points and coefficients can be normalized to remove homothetic, translational and rotational differences between outlines (Rohlf & Archie, 1984; Crampton, 1995; A. J. Haines & Crampton, 2000; Adams, Rohlf & Slice, 2004; Bonhomme *et al.*,

2014). Shape analyses were carried out using the Momocs (“Morphometrics with R”) (Bonhomme *et al.*, 2014) package with the R v4.2.2 software (R Core Team, 2022).

**EFA of outlines: acquisition, processing and analysis**

- High-resolution digital images of lateral and ventral shell views were acquired using a Nikon D3300 camera fitted with a Sigma 105mm f/28 EX DG Macro lens;
- The photographs were processed with Adobe Photoshop, centered, and consistently aligned;
- The photographs were then converted into black masks on a white background (greyscale, 8-bit), and only the shapes of intact shells were retained;
- Outlines were isolated, converted into a list of (*x*; *y*) pixel coordinates, smoothed and used as input data (Supplementary Material Fig. S1).

Outlines were processed before calculating elliptic Fourier transforms following the protocol of Telesca et al. (2018):

- An outline alignment through geometric operations was directly performed on the list of coordinates. This *a priori* normalisation was required to avoid potential bias introduced by the numerical adjustment of shapes prone to bad alignment (usually circular or with bilateral symmetry). Indeed, for oyster outlines, a “first harmonic” normalization approach (J. Claude, 2008) resulted in poor numerical alignment, leading in not homologues elliptic Fourier descriptors;
- Outlines were first smoothed to remove any noise introduced during the digitization process, centred, and outline coordinates were rescaled by their centroid size;
- An equal number of points was sampled along each outline (1,000 pseudo-landmarks);
- Point configurations were aligned through a Procrustes superimposition (Bookstein, 1991; J. Claude, 2008) and starting points were normalised;
- An EFA was then computed on the resulting coordinates from shapes invariant to outline size, rotation and position;
- After preliminary calibration, through inspection of i) the outline reconstruction efficiency and ii) the spectrum of harmonic Fourier power, seven harmonics were identified to encompass 99% of the total harmonic power (Crampton, 1995) (Supplementary Material Fig. S2 A,B);
- Four coefficients per harmonic (a total of 28 descriptors) were extracted for each outline and used as variables quantifying the geometrical information (Rohlf & Archie, 1984; J. Claude, 2008).

The shape information contained in the outlines was then quantified and analysed with classical multivariate tools:

- Principal component analysis (PCA) with a singular value decomposition method without rescaling was performed to summarize the calculated harmonic coefficients in order to define axes capturing the most of the shape variation among individuals (M. Claude, 2013; Telesca *et al.*, 2018);
- The first two shape variables (shape-PC1 and shape-PC2), capturing 83% of the shape variance and describing distinguishable shell outline features, were then used for modelling shell shape variation with environmental gradients (Supplementary Material Fig. S2 C).

SUPPLEMENTARY METHODS S2

**Protocol for Thermal Gravimetric Analyses (TGA)**

Thermogravimetric Analyses are reported following the guidelines made by the Committee on Standardisation of the International Confederation for Thermal Analysis and Calorimetry (ICTAC) and appeared in standards as ASTM E 472 (P. J. Haines, 2002; Gaisford, Kett & Haines, 2016).

**A) Properties of the sample**

i) *Source of material and identification*
- Shell of wild eastern oyster (*Crassostrea virginica*).
- Foliated and chalk microstructure composed of calcite ($CaCO_3$), variable amount of organics (~1-6%) with traces of quartzite ($SiO_2$) and magnesium (Mg).

*ii) Sample history*
- Shells were cleaned, rinsed with milli-Q water, dried at room temperature for seven days.
- The periostracum was removed by sanding, and a tile of prismatic layer isolated (8 × 5 mm) with a Dremel rotary tool.
- Samples were cleaned in an ultrasonic bath (Ultrasonic Cleaner CD-4800, Practical Systems Inc., Odessa, FL, USA) with mill-Q water, air-dried and powdered with an agate mortar.
- All samples were dried in an oven (30 °C for 24 h, convection oven) to remove residual pre-treatment water.

*iii) Physical properties*
- Fine grade powder.

**B) Experimental conditions**

*i) Apparatus used*
- Thermogravimetric Analyser: TGA Q550, TA instrument (New Castle, DE, U.S.A.) Q series.

*ii) Thermal treatment*
- Initial temperature, ~25°C (room temperature).
- Final temperature, 700°C.
- Linear rate of heating, 10°C $min^{-1}$.

*iii) Sample atmosphere*
- Dynamic (flowing) atmosphere.
- Flow rate for balance 40 ml $min^{-1}$ and for sample 60 ml $min^{-1}$.
- Gas composition: nitrogen, “white spot”.

*iv) Sample holder*
- Platinum crucible, cylindrical: diameter 10 mm and height 1.5 mm.
- Sample was tipped and spread to cover the bottom of the crucible.

*v) Sample mass*
- 10 mg of powder were weighted on a separate micro-balance (Ultramicro 4504 MP8, Sartorius, Göttingen; readability 0.1 µg).

**C) Data acquisition and manipulation methods**

*i) Software version*
- Universal Analysis 2000, version 4.5A, TA instrument (New Castle, DE, U.S.A.).
